# Dissecting Gap Junctional and Ephaptic Contributions to Electrical Conduction in a Novel Cardiomyocyte Pair Model

**DOI:** 10.64898/2026.03.05.709926

**Authors:** Xiaobo Wu, Sharon A Swanger, Linnea Elisabeth Bornkast Meier, Clare Dennison, Seth H. Weinberg, Steven Poelzing, Robert G. Gourdie

## Abstract

Electrical communication between excitable cells depends on both direct gap junction (GJ) currents and field-mediated ephaptic interactions, but their relative contributions have remained difficult to quantify experimentally, limiting mechanistic insight into arrhythmia and other disorders of bioelectric signaling in excitable tissues. Building on the concept of a nanoscale, sodium channel-rich perinexus at the cardiac intercalated disc, we developed a ‘Single-on-Paired’ (SoP) preparation in which whole-cell sodium current is recorded from one adult ventricular myocyte that remains end-to-end coupled to an intact partner. This configuration revealed a composite two-cell sodium current characterized by a unique pre-peak waveform which exhibits two slopes in the rising phase, and a pronounced activation jump in sodium current amplitude between closely spaced voltage steps. This feature was absent in isolated single myocytes and interpretable as an ‘Intercalated Disc Signature’ of intercellular activation. By combining graded GJ inhibition, perinexal widening via a Scn1b-derived competitive adhesion peptide, and controlled modulation of extracellular sodium, we show that low sodium conditions favor GJ–dominated activation, whereas at higher, more physiological sodium levels, perinexus-centered ephaptic mechanisms provide substantial support for intercellular activation. A complementary two-cell computational model, simulating our SoP model and incorporating lateral and junctional sodium channels, reproduces the two-cell ‘Intercalated Disc Signature’ and predicts a shift from GJ-dominated activation at low sodium to ephaptic-dominated support at higher, more physiological sodium concentrations when GJ conductance is reduced. Together, these results provide direct experimental evidence for a specific structural and molecular substrate of ephaptic coupling in the heart and establish a framework for dissecting how nanoscale extracellular cleft geometry, channel organization, sodium level, and GJ conductance jointly tune electric-field-based mechanisms of activation in excitable tissues, with implications for paradoxical clinical responses to anti-arrhythmic interventions.

## INTRODUCTION

Electrical communication between excitable cells underlies phenomena as diverse as rhythm generation and action-potential propagation in the heart and the emergence of large-scale activity in the brain, yet the relative roles of direct and electric-field-mediated coupling remain incompletely defined.^1–3^ In both tissues, multiple mechanisms - including chemical synaptic transmission, gap junctional coupling (GJC), and ephaptic coupling (EpC) - contribute to intercellular electrical signaling. Traditionally, GJC, mediated by gap junctions (GJs) - clusters of connexin channels that provide direct cytoplasmic continuity - has been recognized as an important, but not exclusive, mechanism of electrical communication in the brain^4,5^ and the principal means of intercellular electrical transmission in the heart.^6^ In contrast, EpC, mediated by extracellular electric fields between closely apposed membranes, remains under active investigation in both the brain and heart.^7,8^

In the brain, EpC has been well-recognized^9,10^ and field effects in confined extracellular spaces have been implicated in modulation of interneuronal synchrony, epileptiform activity, and candidate mechanisms for memory-ensemble formation and conscious processing.^11–15^ EpC in the heart also has a long history,^16,17^ and emerging experimental and computational studies increasingly support the view that it contributes to electrical conduction ^18–22^ and can participate in arrhythmogenesis ^18,23,24^. Nevertheless, direct measurement of EpC between cardiomyocytes is still required to confirm its existence and quantify its contribution to conduction. Classic connexin-43 (Cx43) loss-of-function studies show that cardiac conduction can persist when GJC is markedly reduced,^25–28^ suggesting that additional, potentially ephaptic mechanisms maintain propagation, but the relative contributions of residual direct coupling versus field effects remain unresolved.

EpC depends on extracellular electric fields, and it is proposed to occur within the narrow intercellular cleft between apposing cells.^21,29,30^ Subsequent work has identified such a nanoscale cleft within the cardiac intercalated disc (ID), the perinexus - a region enriched with voltage-gated sodium channels (Nav1.5) and located adjacent to Cx43-formed GJs.^3,18,31–33^ Experimental and computational evidence from multiple groups support a model in which rapid activation of sodium channels within the perinexus lowers the local extracellular potential to trigger EpC, thereby trans-activating sodium channels in neighboring cardiomyocytes. ^19,20,22,34^

Despite extensive investigation of GJC and EpC in cardiac conduction, a fundamental question remains unresolved: do they operate as independent or cooperative mechanisms? Current understanding relies heavily on computational models with inherent limitations,^21,35–40^ and experimental efforts have been hampered by the difficulty of isolating EpC from GJC. For example, Cx43 knockout or loss-of-function studies demonstrate that the conduction of electrical impulses can persist,^41,42^ yet controversy continues over the extent to which residual GJC shapes the observed propagation of action potentials, complicating attempts to ascribe conduction properties specifically to EpC.

A central obstacle has been the lack of experimental systems that preserve native IDs while allowing direct, quantitative dissection of junctional and field-mediated components of activation at the level of isolated cell pairs. Here we introduce a ‘Single-on-Paired’ (SoP) cardiomyocyte preparation that addresses this gap. In this model, whole-cell sodium current is recorded from one adult ventricular myocyte that remains end-to-end coupled to an intact partner via a native ID containing both Cx43 gap-junction plaques and perinexal clefts, revealing a distinctive “Intercalated Disc Signature” that reports intercellular activation. By combining graded GJ inhibition, perinexal widening with a Scn1b-derived competitive adhesion peptide, controlled modulation of extracellular sodium, and a complementary two-cell computational model simulating the SoP experiment and incorporating both lateral and junctional sodium channels, we dissect how GJC and perinexus-centered EpC jointly support activation under different ionic and structural conditions.

Beyond clarifying how gap-junctional and ephaptic mechanisms cooperate to preserve cardiac conduction when Cx43 conductance is compromised, this SoP framework establishes an experiment-ready paradigm for probing Cx43-anchored nanodomains in excitable tissues. Given the presence of structurally analogous Cx43 networks, voltage-gated sodium channels, and nanoscale clefts in astrocyte syncytia and other Cx43-rich systems,^2,43^ the mechanisms uncovered here may have broad relevance for understanding how local microdomains of coupled cells shape bioelectric field signaling from the nanoscale to organ-level dynamics in both heart and brain.

## METHODS

### Cardiomyocyte isolation

Ventricular cardiomyocytes from adult male and female C57BL/6 mice (8–12 weeks) were isolated using a Langendorff-free protocol^44^ modified to increase the yield of cardiomyocyte pairs with preserved intercalated discs. Animals were anesthetized with isoflurane and euthanized by cervical dislocation. The chest was opened, the heart exposed, and the abdominal aorta was transected for right-ventricular perfusion. A 27.5-G needle attached to a syringe containing perfusion buffer (in mM: 130 NaCl, 5 KCl, 0.5 Na_2_HPO_4_, 10 HEPES, 1 MgCl_2_,10 taurine, 10 glucose, and 25 blebbistatin) was inserted into the right ventricle near the apex to flush blood from the heart (∼2 ml over 1 minute). The heart was then excised and placed in a dish. The aorta was clamped, and the same needle was advanced through the apical hole into the left ventricle to deliver an additional 4 ml of perfusion buffer over 2 minutes, clearing blood from the ventricles and coronary arteries. When effluent was free of blood, the heart was transferred to a second dish containing 3 ml of digestion solution (perfusion buffer plus 550 U/ml collagenase II, Worthington-Biochem, USA) pre-warmed to 37°C. Using the same apical access, the left ventricle was perfused with 20 ml digestion solution over 10 minutes on a 37°C heating pad, producing modest digestion that reproducibly yielded usable numbers of myocyte pairs. Atria were removed and the ventricles transferred to a fresh dish containing 3 ml of digestion solution, mechanically separated, and gently agitated for 2 minutes. Five ml of stop solution (40 ml perfusion buffer plus 4 ml fetal bovine serum) were then added, followed by an additional 2-minute agitation. Remaining tissue chunks were removed, and the cell suspension was transferred to a 50-ml conical tube and centrifuged, ramping to 1000 rpm at room temperature. The supernatant was discarded and the pellet resuspended in stop solution to 10 ml; this wash was repeated once more before calcium reintroduction. A 100 mM CaCl_2_ stock was then added to the suspension in three steps (30, 30, and 40 µl at 30-s intervals) to reach a final Ca^2+.^ concentration of 1 mM.

### Patch clamp

Whole-cell sodium current recordings from single and paired cardiomyocytes were performed at room temperature in a voltage-clamp mode. Recordings were obtained with an Axopatch 200B amplifier, Digidata 1550B digitizer, and pClamp 11 software (Molecular Devices). Patch pipettes had a tip resistance of 2.5–3 MΩ, and series resistance was compensated by up to 70%. Cell capacitance in single and paired cardiomyocytes was measured at a holding potential of +10 mV. The voltage-clamp protocol consisted of a holding potential of −100 mV for 2 s followed by 300-ms steps from −80 mV to +20 mV in 5-mV increments. Only paired cardiomyocytes attached end-to-end with clearly visible intercalated discs by phase contrast microscopy were selected for SoP recordings. The internal (pipette) solution for whole-cell sodium-current recordings contained (in mM): 120 cesium methane sulfonate, 20 tetraethylammonium chloride, 2 MgCl₂, 10 HEPES, 10 EGTA, and 4 adenosine 5′-triphosphate magnesium salt, adjusted to pH 7.3 with cesium hydroxide. The external solution contained (in mM): 25 NaCl, 120 N-methyl-D-glucamine, 10 HEPES, 1.8 MgCl₂, 1.8 CaCl₂, and 10 glucose, adjusted to pH 7.3 with HCl. Verapamil (100 µM) was added to the external solution to block calcium channels.

### Fluorescent Immunolabeling

Isolated cardiomyocytes were plated on laminin-coated slips for 2 hours before fixation with 4% PFA for 15 minutes. Cardiomyocytes were then rinsed with 1x PBS for 3 times with 5 minutes per rinse. A 0.1% Triton X-100 solution was added on each slip for 5 minutes followed by the rinsing with 1x PBS for 3 times with 5 minutes per rinse. The 1x BSA blocking solution was added on each slip for 30 mins. Cardiomyocytes were labeled with a rabbit polyclonal antibody against either sodium channel Na_v_1.5 (1:250, S0819, MilliporeSigma) or β1/β1b (1:200, Epitope: 44KRRSETTAETFTEWTFR60^18,45^ alongside with a mouse monoclonal antibody against Cx43 (1:250, sc-271837, Santa Cruz Biotechnology) and incubated at 4°C overnight. After a rinsing off primary antibody (with 1x PBS), cardiomyocytes were then second labeled with a donkey anti-rabbit AlexaFluor 568 (1:2000, ThermoFisher Scientific) and a donkey anti-mouse AlexaFluor 647 (1:2000, ThermoFisher Scientific) for 1.5 hours. After rinsing with 1x PBS, nuclei were stained with Hoescht 33342 for 10 minutes (1:30,000, Invitrogen), rinsed in 1x PBS again and slides were then cover-slipped for imaging, as we have described previously^18^.

### Confocal microscopy

Confocal microscopy was performed with a TCS SP8 laser scanning confocal microscope equipped with a Plan Apochromat 63×/1.4 numerical aperture oil immersion objective and a Leica HyD hybrid detector (Leica). Imaging each fluorophore was performed sequentially, and the excitation wavelength was switched at the end of each frame.

### Transmission electron microscopy

Isolated cardiomyocytes were treated either with 50-µM scr1 or βadp1 for 1 hour before fixation with 2% glutaraldehyde. Cardiomyocyte suspensions were pelleted by centrifuge and post-fixed with 2% osmium tetroxide for 1 hour on ice, followed by en bloc staining with 1% uranyl acetate in 25% ethanol for 1 hour. Samples were then dehydrated through a graded ethanol series (50%, 70%, 90%, and 100%, with two changes at 100%). Embed 812 resin (EMS #14900) was infused overnight using a 1:1 mixture of propylene oxide and Embed 812, followed by four changes of 100% resin. Samples were embedded in BEEM capsules, and resin was polymerized at 65 °C for 48 hours. Ultrathin sections (70 nm) were cut using a Leica EM UC7 ultramicrotome and collected on gold grids. Imaging of the perinexus between paired cardiomyocytes was performed at x13,000 magnification using a FEI tecnai T12 transmission electron microscope. For each perinexus, the perinexal width (W*_p_*) was calculated over the distance of 50 nm from the edge of the gap junction plaque using a previously reported method.^46^

### Two-cell computational modeling

A two-cell computational model was established to elucidate the ‘Intercalated Disc Signature’ (IDS) in sodium current traces and the influence of extracellular sodium levels on intercellular conduction between two cells. Briefly, this model simulates two myocytes electrically coupled via end-to-end intercalated discs, and it is assumed that sodium channels in each myocyte were distributed on the lateral and junctional membranes, with the junctional membrane adjacent to the cleft (comparable to the perinexus). The cell model also incorporates direct electrical coupling via gap junctions (Figure 6, top). The model is based on a previously implemented model of ephaptic-mediated automaticity in cell pairs^24,47^ and an electrical circuit representing the two coupled cells is shown in Figure 6 (bottom).

In this model, there are two intracellular potentials *V_1_* and *V_2_* in each cell (assuming the bulk extracellular potential is electrically grounded), and a single voltage in the intercellular cleft V_cl_ representing the cleft potential. *C_lat_* and *C_junc_* respectively denote the lateral and junctional membrane capacitances. *C_lat_* = *C_m_A_lat_*, where *C_m_* is the unit membrane capacitance and *A_lat_* = *2𝜋rL* is the lateral membrane area (*r* is the radius of the cross section and *L* is the length of the cell); *C_junc_* = *C_m_A_junc_*, where *A_junc_* = *𝜋r^2^*.

To simulate the compartmental sodium currents, a voltage step clamp protocol was applied to this two-cell model by delivering a series of voltage steps to a node inside Cell 1 labeled as V_1_. Full model equations are provided in our prior work,^24^ with the modification that V_1_ dynamics are clamped to a fixed voltage. Ionic current dynamics are represented by the Luo-Rudy 1991 model.^48^ Importantly, note that the cleft (simulated perinexus) width modulates electrical dynamics in two ways: 1) cleft width is inversely proportional cleft resistance (R_cl_ in Figure 6), such that a wider cleft has a lower cleft resistance, and 2) cleft width is proportional to cleft volume, which impacts cleft ionic concentration changes, such that a given junctional current modulates cleft concentration to a greater extent for a smaller cleft volume.

We simulate the two-cell model dynamics, varying the cleft widths, gap junction conductance, and bulk extracellular sodium concentrations in a given simulation for fixed voltage clamp, and measure the sodium currents from each compartment of a cell and the cleft potential.

**Table 1.**
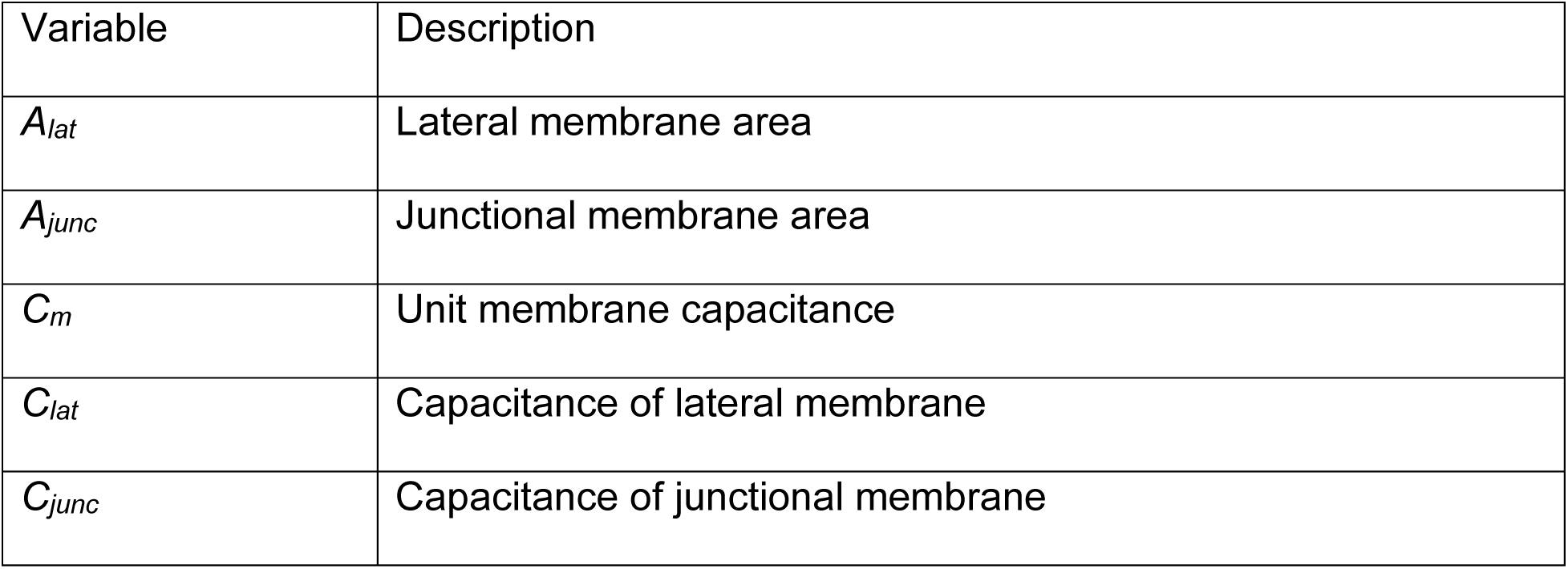
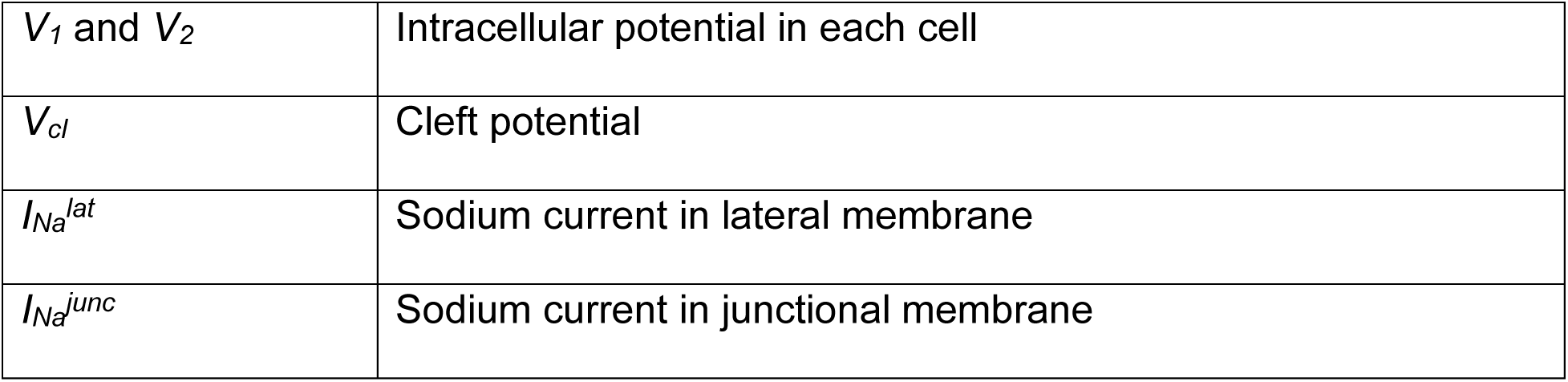
Description of model parameters.

**Table 2.** Parameter values.

| Parameter | Value (units) |
| --- | --- |
| $r$ | 11 ($\mu\text{m}$) |
| $L$ | 100 ($\mu\text{m}$) |
| $A_{lat}$ | $2\pi rL$ ($\mu\text{m}^2$) |
| $A_{junc}$ | $\pi r^2$ ($\mu\text{m}^2$) |
| $C_m$ | $1 \times 10^{-8}$ ($\mu\text{F}/\mu\text{m}^2$) |
| $C_{lat}$ | $C_m A_{lat}$ ($\mu\text{F}$) |
| $C_{junc}$ | $C_m A_{junc}$ ($\mu\text{F}$) |
| $[\text{Na}^+]_o$ | 25 or 145 (mM) |
| $[\text{K}^+]_o$ | 5.4 (mM) |
| $[\text{Ca}^{2+}]_o$ | 1.8 (mM) |
| F | 96.5 (C/mM) |
| R | 8.314 (J/K) |
| T | 310 (K) |

### Statistical Analysis

All data are presented as mean ± SEM unless otherwise stated. Statistical analyses were performed using GraphPad Prism (Version 10.5, GraphPad Software, San Diego, CA). For comparisons between two groups, unpaired or paired two-tailed Student’s *t*-tests were used as appropriate. For comparisons involving more than two groups, one-way analysis of variance (ANOVA) was performed, followed by Tukey’s post hoc multiple-comparison tests. A *p* value < 0.05 was considered statistically significant.

## RESULTS

### Establishment and Characterization of the ‘Single-on-Paired’ Experimental Model

Dissecting the individual contributions of gap junctional coupling (GJC) and ephaptic coupling (EpC) at the cellular level is essential for understanding the mechanisms governing electrical conduction between cardiomyocytes. While previous cardiomyocyte preparations have provided important insights, none have directly enabled controlled interrogation of electrical coupling mechanisms at the cellular scale. To address this limitation, we developed a novel ‘Single-on-Paired’ (SoP) experimental model that applied the single patch-clamp configuration to paired, end-to-end connected cardiomyocytes that retained an intact intercalated disc (ID), including gap junction (GJ) and perinexal domains. Myocyte pairs displaying extensive side-by-side contact along their longitudinal axes were excluded to avoid potential confounding effects of non-disc lateral coupling.

Within this end-to-end model, the whole-cell sodium current (I_Na_) of paired cardiomyocytes was measured to evaluate electrical conduction, because activation of voltage-gated sodium channels (Nav1.5) is critical for both GJC and EpC. Theoretical considerations indicate that widening perinexal space in paired cardiomyocytes should alter local I_Na_ within this restricted cleft while leaving non-cleft regions relatively unaffected (Figure 1a). Therefore, we hypothesized that increasing perinexal width would augment the whole-cell I_Na_ of paired cardiomyocytes.

To test this hypothesis, we first validated the feasibility of the SoP model for measuring whole-cell I_Na_ from both cardiomyocytes. Single and paired cardiomyocytes were isolated from adult mouse ventricular tissues. Single patch-clamp recordings in the voltage-clamp mode were then performed at room temperature using an extracellular sodium concentration of 25 mM - a reduced sodium level commonly employed to facilitate stable measurement of voltage-gated I_Na_ in cardiac myocytes.

**Figure 1.**
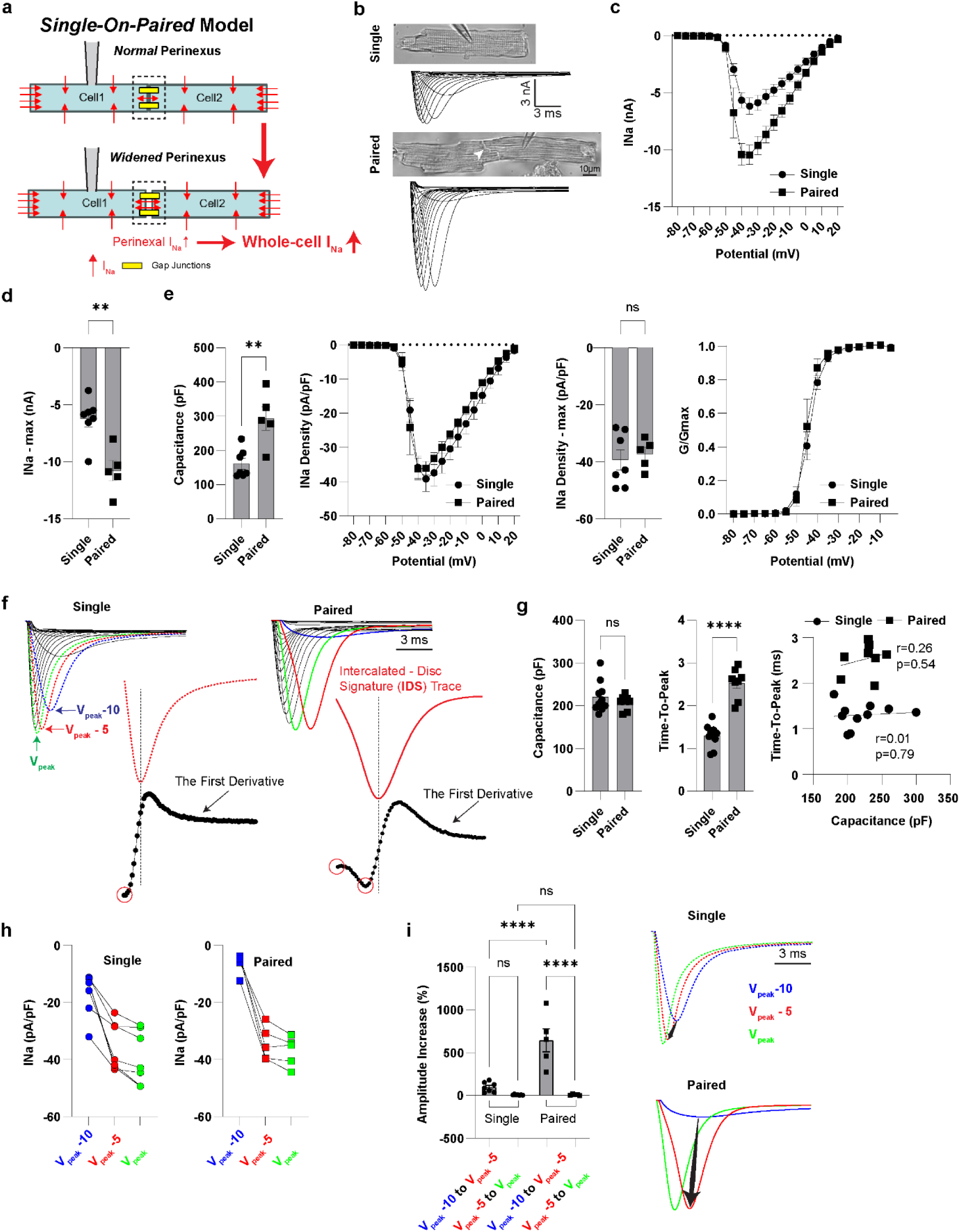
Establishment and characterization of the *‘Single-on-Paired’* cell model. **a.** Rationale of the ‘Single-on-Paired’ (SoP) model. **b**. Representative whole-cell I_Na_ traces from single and paired cardiomyocytes. **c**. Peak I_Na_ I-V curves from single (n=7) and paired cardiomyocytes (n=5). **d**. Max I_Na_. **e**. Capacitance, I_Na_ density I-V curve, max I_Na_ density and steady-state activation. **f**. Normalized I_Na_ traces from single and paired cardiomyocytes with the pre-peak trace (red), whose characteristics define the Intercalated Disc Signature (IDS), followed by its first derivative. **g**. Capacitances and time-to-peak duration from *selected* single (n=9) and paired cardiomyocytes (n=8), and the correlation between capacitance and time-to-peak duration. **h**. I_Na_ densities from the three consecutive traces at V_peak_-10, V_peak_-5 and V_peak_ from single and paired cardiomyocytes described in Figure 1c-e. **i***. Left*: The increase in amplitude from V_peak_-10 to V_peak_-5 and V_peak_-5 to V_peak_ respectively; *Right*: The representative 3 consecutive traces at V_peak_-10, V_peak_-5 and V_peak_ from single and paired cardiomyocytes (black arrows indicate the increase in amplitude from V_peak_-10 to V_peak_-5). Unpaired *t*-test was used for **d**, **e**, and **g**, Pearson correlation test was used for **g** and One-way ANOVA followed by Tukey correction for multiple comparison for **i**: \**p*<0.05;\*\**p*<0.001;\*\*\**p*<0.0001;\*\*\*\**p*<0.00001; ns, not significant.

It was found that in contrast to single cardiomyocytes, paired cardiomyocytes exhibited larger whole-cell I_Na_ amplitudes (Figure 1b). To characterize the properties of whole-cell I_Na_ in paired cardiomyocytes, we first employed a conventional approach by comparing I_Na_ amplitudes between single and paired cardiomyocytes. At first, peak I_Na_ current-voltage (I-V) relations were quantified, and it was found that compared with single cardiomyocytes, paired cardiomyocytes exhibited large amplitudes at multiple voltage steps with the largest one at -10.80 ± 0.88 nA relative to -6.22 ± 0.71 nA in single cardiomyocytes (Figure 1b, 1c and 1d). In addition, compared with single cardiomyocytes, estimated capacitance in paired cardiomyocytes was nearly doubled (single: 160.7 ± 15.74 vs paired: 293.2 ± 34.86 pF; Figure 1e), resulting in comparable I_Na_ densities to single cardiomyocytes (Max I_Na_ density: -39.34 ± 3.53 pA/pF for single and -37.29 ± 2.33 pA/pF for paired, Figure 1e). Steady-state activation was also similar between single and paired cardiomyocytes (V_mid_: -44.04 ± 0.93 mV for single; -45.01 ± 1.26 mV for paired, Figure 1e).

Although conventional comparisons showed similarities in I_Na_ properties between single and paired cardiomyocytes, closer inspection of normalized I_Na_ traces in both cell types (Figure 1f), revealed interesting and important differences in the morphology of traces preceding peak activation. Specifically, compared with single cardiomyocytes, the peak I_Na_ trace (green) and its two preceding traces (red and blue) in paired cardiomyocytes displayed a characteristic progression. For ease of interpretation, we used V_peak_ to denote the voltage step for the peak I_Na_ trace, V_peak_−5 (5 mV below V_peak_) for the preceding red trace, which we also refer to as the pre-peak trace, and V_peak_−10 (a further 5-mV step down) for the blue trace. The first notable feature of this three-trace sequence in paired cardiomyocytes was a large, non-linear increase in amplitude between the V_peak_−10 and pre-peak V_peak_−5 traces that was consistently present, whereas single cardiomyocytes showed a more linear, gradual change in amplitude across the same 5-mV steps, with no abrupt change in apparent channel activity. We hereafter use the shorthand ‘activation jump’ to refer to this unique feature of the pre-peak I_Na_ (i.e. V_peak_−5) trace in paired cardiomyocytes.

The second distinct feature of the three-trace sequence in paired cardiomyocytes was that the activation phase of the pre-peak curve reliably exhibited a *two-slope* morphology, while the bracketing V_peak_ and V_peak_−10 traces, as well as all other voltage steps, displayed only a *single* activation slope. Similarly, in single cardiomyocytes, all traces showed a *single-slope* activation. When a first-derivative plot of the pre-peak trace was generated, this phenomenon became particularly clear, with the two-slope activation unique to paired cardiomyocytes appearing as two distinct troughs.

Across all analyzed recordings, the three--trace sequence and its associated pre-peak (i.e. V_peak_−5) amplitude jump and two-slope activation pattern were observed only in bona fide paired cardiomyocytes and never in single ones. As an additional empirical control, further I_Na_ recordings were obtained from large single cardiomyocytes, as indicated by capacitance (>180 pF), and compared to paired cardiomyocytes with similar capacitances (Figure 1g, left). In paired cells, the two-slope trace at V_peak_−5 displayed a markedly prolonged time-to-peak during the rising phase relative to the corresponding trace from single cells (Figure 1g, middle), which showed only single-slope activation (trace not shown), indistinguishable from single cells with capacitances <180 pF. Moreover, plots of capacitance versus time-to-peak revealed no meaningful correlation in any cell group (Figure 1g, right). Together, these observations indicate that under comparable recording conditions and cell sizes, the IDS morphology depends strictly on the presence of an intercalated disc between two cells and is not reproduced in isolated single cardiomyocytes, large or otherwise. In this configuration, the controlled loss of ideal voltage control in the downstream cell and cleft serves as the biophysical signal that we interpret as reporting activation across the intercalated disc.

Next, I_Na_ densities were measured for the three traces at V_peak_-10, V_peak_-5 and V_peak_ in single (those in Figure 1e) and paired cardiomyocytes (Figure 1h), and the relative changes in amplitude were quantified to assess the magnitude of the activation jump (Figure 1i, *left*). Compared with single cardiomyocytes, paired cells showed a pronounced and highly significant increase in amplitude from V_peak_-10 to V_peak_-5, but only a modest additional increase between V_peak_-5 and V_peak._ The activation jump from V_peak_-10 to V_peak_-5 in paired cardiomyocytes is illustrated in Figure 1i-*right*. This pattern suggested that the two-slope V_peak_-5 trace in paired cardiomyocytes reflects, in sequence, first the well-established dyssynchronous recruitment of I_Na_ in the patched cell, and then unclamped activation of sodium channels in the downstream, unpatched cell.

### Characterization of the Intercalated-Disc-Signature and its role in electrical coupling

The two-slope waveform and activation jump of the V_peak_-5 pre-peak trace observed in paired cardiomyocytes is referred to as the Intercalated Disc Signature (IDS). The IDS was associated with a significantly longer time-to-peak at V_peak_-5 in paired cardiomyocytes than in single cardiomyocytes (Figure 2a). By contrast, the decay time, reflecting sodium channel inactivation, was markedly shorter in paired as compared to single cardiomyocytes (Figure 2b). Notably, whilst clear twin slopes were never discerned for the V_peak_ curve following the two-slope V_peak_-5 trace (Figure 2c), the prolongation of time-to-peak during the rising phase and the shortening of decay time during the decay phase for V_peak_ remained comparable to that seen for the V_peak_-5 curve (Figure 2d).

Because the IDS waveform appears only in cell pairs attached by an ID, we interpreted its presence as an operational readout of trans-junctional intercellular activation across the intercalated disc unique to our voltage-clamp SoP configuration. We therefore use the term ‘trans-junctional’ here to denote activation across the intercalated disc in this voltage-clamp configuration, rather than classical current-clamp propagation between two unclamped cells.

To test whether the IDS can be used to evaluate the trans-junctional biophysics of intercalated disc-attached cardiomyocytes, we applied GJ inhibitors, including 2-aminoethoxydiphenyl borate (2-APB)^49,50^ to paired cardiomyocytes and assessed whether inhibition prolonged the IDS time-to-peak. Although several GJ inhibitors (carbenoxolone, 18β-glycyrrhetinic acid, and 2-APB) were examined, cardiomyocytes only remained viable over 60 minutes with 2-APB treatment.

**Figure 2.**
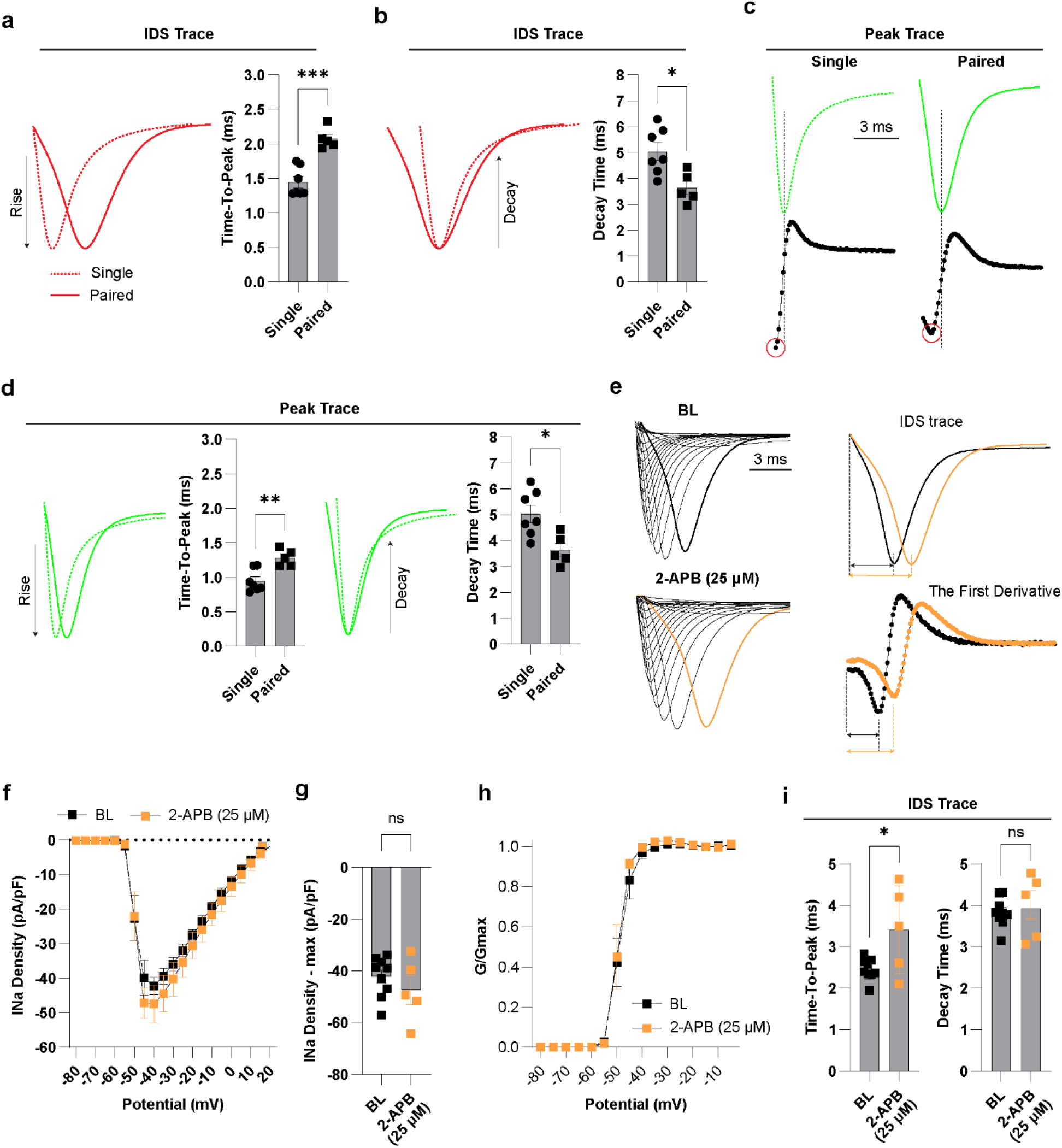
Establishment and characterization of the *‘Single-on-Paired* model (*continued*). **a**. Time-to-peak for the rising phase of the IDS. **b**. Decay time for the decay phase of the IDS. **c**. Representative peak traces and their first derivative from single and paired cardiomyocytes described in Figure 1f. **d**. The time-to-peak and decay time of peak trace from single and paired cardiomyocytes described in Figure 1f. **e**. Representative normalized whole-cell I_Na_ at BL (n=9) and after 2-APB (25 µM) treatment (n=5) in paired cardiomyocytes and the IDS with its first derivative. **f**. INa densities at BL and after 2-APB (25 µM) treatment in paired cardiomyocytes. **g**. Max INa densities at BL and after 2-APB (25 µM) treatment. **h**. Steady-state activation at BL and after 2-APB (25 µM) treatment. **i**. The quantified time-to-peak and decay time at BL and after 2-APB (25 µM) treatment in paired cardiomyocytes. Unpaired t-test was used for statistical analysis for **a**, **b**, **d**, **f**, **g** and **i**. \**p*<0.05;\*\**p*<0.001;\*\*\**p*<0.0001; ns, not significant.

Paired cardiomyocytes were treated with 25-μM 2-APB for at least 30 min before recording. Although 2-APB did not significantly alter I_Na_ properties relative to baseline (BL) (Figure 2f-h), it markedly prolonged the IDS rising phase (Figure 2e and 2i; BL: 2.46 ± 0.09 ms vs 25-μM 2-APB: 3.41 ± 0.47 ms), whereas the decay phase was unchanged (BL: 3.83 ± 0.12 ms vs 25 μM 2-APB: 3.93 ± 0.34 ms). These findings indicate that the IDS trace is sensitive to GJ uncoupling at 25 mM sodium and can be used as a readout of intercellular conduction of activation between cardiomyocytes in this model.

Taken together, these results demonstrate that I_Na_ transients in the SoP model differ substantially from those in isolated single cells, and that the timing of sodium-channel activation within the IDS waveform is sensitive to gap-junctional coupling, at least at the non-physiologic 25 mM concentration of extracellular sodium tested thus far.

### Perinexal remodeling modulates the ‘Single-on-Paired’ sodium current

Next, we asked whether widening the perinexus - our putative EpC substrate - alters whole-cell I_Na_ in the SoP model. Based on the computational predictions, ^20,37,51^ and the concept that perinexal geometry regulates perinexal I_Na_ thus contributing to the net current of paired cardiomyocytes, we hypothesized that under our 25-mM extracellular sodium conditions, perinexal widening would increase whole-cell I_Na_ in SoP cell pairs.

Based on prior work showing that Scn1b β1/β1b adhesion modulates perinexal width and conduction in a manner consistent with EpC,^18,23,52^ we perfused 50 μM of the Scn1b competitive adhesion disruptor βadp1 or its scrambled control peptide (scr1). Treatment with βadp1 for 30 min significantly increased peak I_Na_ density compared with scr1 and BL (Figure 3a; βadp1: −52.21 ± 3.19 pA/pF, scr1: −40.49 ± 2.19 pA/pF, BL: −40.56 ± 2.83 pA/pF), without altering steady-state activation (V_mid_: −48.44 ± 1.27 mV for βadp1, −48.00 ± 1.39 mV for scr1, −48.86 ± 1.10 mV for BL). The increase in I_Na_ was confined to V_peak_, indicating a modest effect under these conditions. Time-to-peak and decay of the IDS waveform were unchanged after βadp1 (Figure 3b). Importantly, the time-to-peak of the peak I_Na_ traces were significantly shortened relative to scr1 and BL, with no change in decay (Figure 3c), consistent with computational predictions of EpC that widening clefts can both reduce I_Na_ self-attenuation (i.e. increase I_Na_) while simultaneously increasing conduction velocity (i.e. reduced intercellular latency).^21^ As expected, βadp1 did not alter I_Na_ properties in single cardiomyocytes (Figure S1a,b). Together, these data indicate that βadp1 accelerates sodium-channel recruitment and increases peak I_Na_ during supra-threshold activation in paired cells, consistent with an ephaptic contribution mediated by perinexal remodeling.

**Figure 3.**
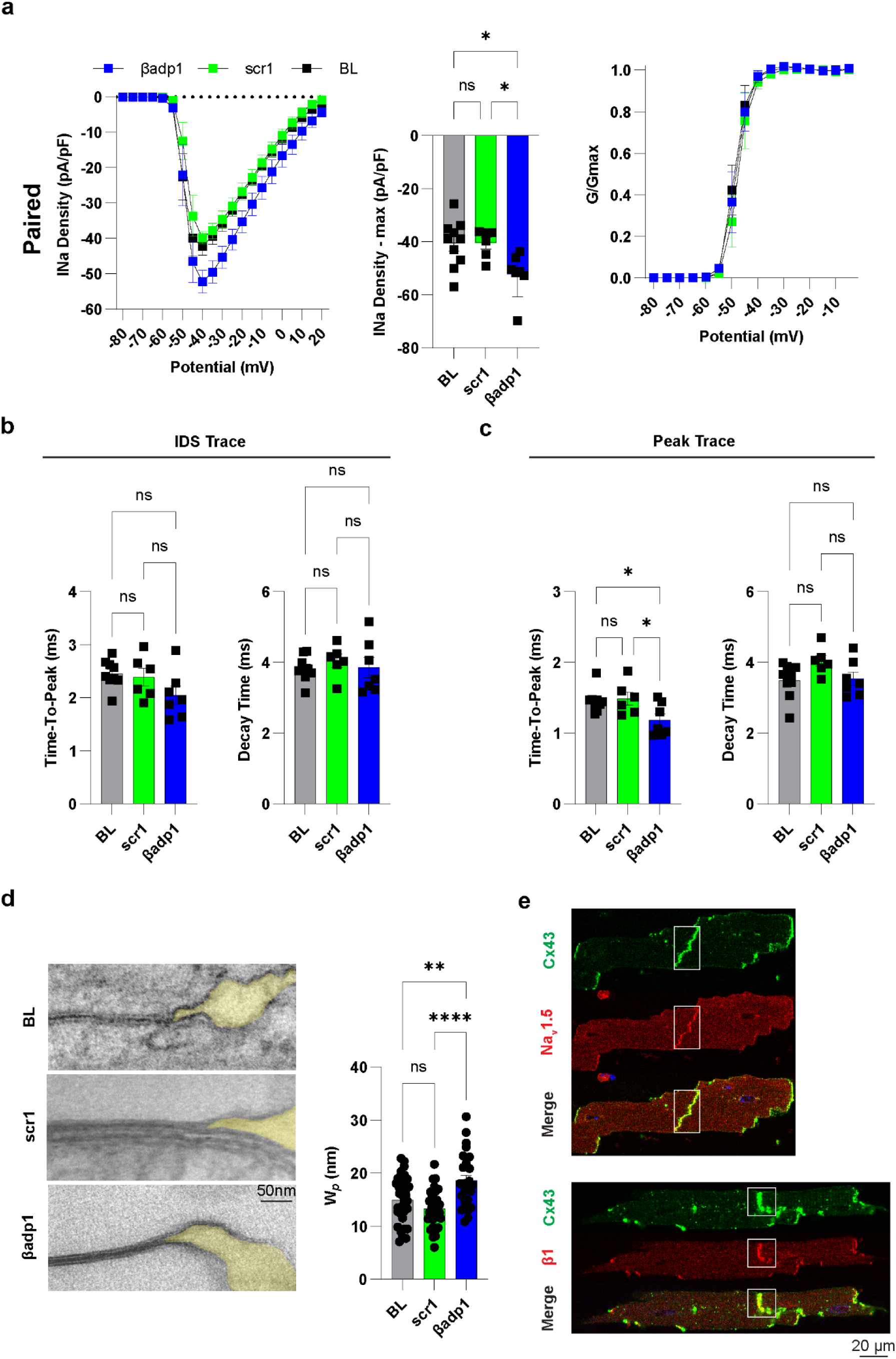
Perinexal remodeling increases sodium current in paired cardiomyocytes. **a**. I_Na_ density I-V curves, max I_Na_ densities and steady-state activation curves at BL (n=10), scrambled βadp1 (scr1, n=6) and βadp1 (n=7) in paired cardiomyocytes. **b**. Time-to-peak and decay time of the IDS traces at BL, scr1 and βadp1. **c**. Time-to-peak and decay time of the peak traces at BL, scr1 and βadp1. **d**. Representative images of perinexus in paired cardiomyocytes and the quantified perinexal width (W*_p_*) at BL (perinexi: n=32), scr1 (perinexi: n=31) and βadp1 (perinexi: n=27). **e**. Co-localization of Cx43, Nav1.5 and β1/β1b within IDs between paired cardiomyocytes. One-way ANOVA followed by Tukey correction for multiple comparison. \**p*<0.05;\*\**p*<0.001;\*\*\*\**p*<0.00001; ns, not significant.

To further test whether perinexal widening is associated with increased SoP I_Na_, paired cardiomyocytes were fixed after each intervention and perinexal width (W_p_) was measured by transmission electron microscopy. βadp1 significantly increased W_p_ within the 0–50 nm region immediately adjacent to GJs (Figure 3d; βadp1: 18.65 ± 0.98 nm, scr1: 13.33 ± 0.65 nm, BL: 14.97 ± 0.79 nm). Overall, these findings indicate that local widening of the proximal, near-junctional perinexus in paired cardiomyocytes is accompanied by increased I_Na_ in paired cardiomyocytes and that these observations are consistent with an ephaptic contribution to cell-to-cell coupling in the SoP model.

### Intercalated disc structures in paired cardiomyocytes

We also confirmed the localization of Connexin-43 (Cx43), Nav1.5, and voltage-gated sodium channel β1/β1b subunits at IDs in paired cardiomyocytes (Figure 3e). Although not central to our primary aims, we noted that myocyte isolation induced internalization of both GJs and their adjacent perinexal domains (Figure S1c) - a phenomenon previously reported for GJs before the perinexus was defined.^53^ Consistent with this, isolated cells showed intracellular co-localization of Cx43 with Nav1.5 and β1 subunits (Figure S1d), suggesting that endocytic removal of the ID includes the perinexal compartment. These findings suggest that the isolation procedure can remove Nav1.5, together with GJs and perinexal ion-channel complexes, from remodeled disc regions. This selective loss of channels within the ID may contribute to variability in whole-cell I_Na_,^54^ particularly in isolated single cardiomyocytes, underscoring the value of the SoP preparation, which presumably may better preserve junctional channel complexes.

### Gap Junction Inhibition Abolishes Electrical Conduction Between Cardiomyocytes at Low Extracellular Sodium

As shown above, 25-μM 2-APB delayed trans-junctional sodium-channel activation in the IDS waveform. To test whether stronger GJ inhibition suppresses activation of the unclamped (‘downstream’) cell in the SoP model, we increased 2-APB to 50 μM. Unexpectedly, after ≥30 min treatment, the SoP I_Na_ was reduced by ∼50% compared with BL (Figure 4a,b; 50-μM 2-APB: −4.93 ± 0.35 nA vs BL: −9.70 ± 0.46 nA). Importantly, the IDS (red), which reports trans-junctional activation at BL, was completely absent after 50-μM 2-APB. Of particular note, I_Na_ traces under 2-APB in paired cells were nearly identical to those from isolated single cardiomyocytes, with similar I–V relationships and peak I_Na_, indicating a lack of detectable current in the downstream cell. Together, these observations are consistent with loss of both gap-junctional and ephaptic intercellular activation.

**Figure 4.**
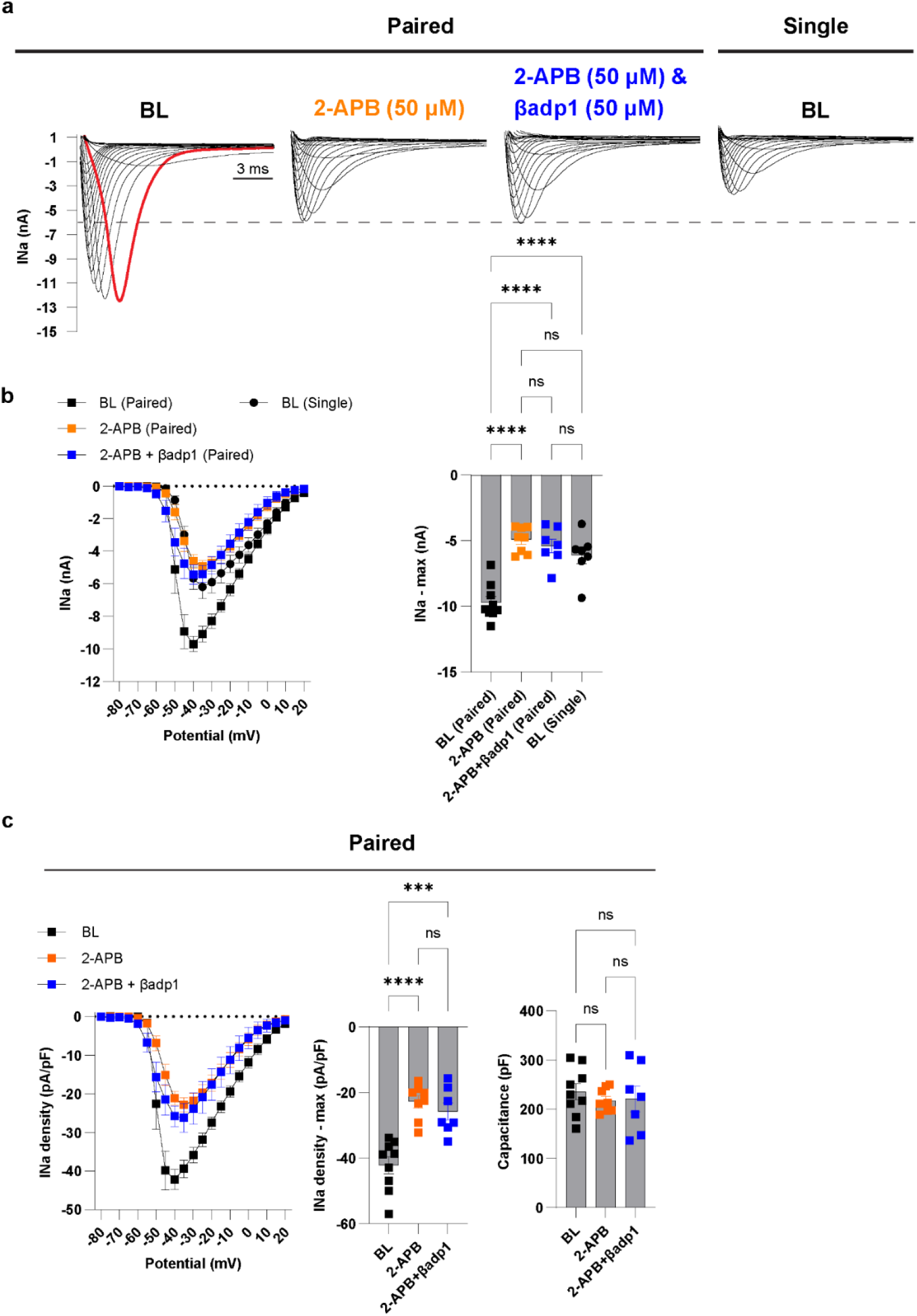
High gap junction inhibition induces electrical conduction block between cardiomyocytes in the low sodium condition. **a**. Representative whole-cell I_Na_ traces at BL, with 50-µM 2-APB and with combined treatments of 50-µM 2-APB and 50-µM βadp1 in paired cardiomyocytes, as well as at BL in single cardiomyocytes (n=7) at 25 mM extracellular Na^+^. **b**. Peak I_Na_ I-V curves and max I_Na_ at BL (n=9), 2-APB (n=8) and 2-APB&βadp1 (n=7) in paired cardiomyocytes as well as at BL in single cardiomyocytes. c. I_Na_ density I-V curves, max I_Na_ densities and capacitance at BL, 2-APB and 2-APB& βadp1 in paired cardiomyocytes. One-way ANOVA followed by Tukey correction for multiple comparison. \**p*<0.05;\*\**p*<0.001;\*\*\**p*<0.0001;\*\*\*\**p*<0.00001; ns, not significant.

Importantly, adding βadp1 under these conditions did not change I_Na_ morphology or magnitude, indicating that perinexal widening cannot rescue electrical conduction when GJs are strongly inhibited and extracellular sodium is very low. Consistent with this, I_Na_ density–voltage relationships and peak I_Na_ density were markedly reduced by 2-APB, either alone or combined with βadp1, relative to BL (Figure 4c, *left* and *middle*). Collectively, these results, together with our finding that perinexal widening augments I_Na_ in paired cardiomyocytes (Figure 3), suggest that EpC is insufficient to support conduction independent of GJC at the low, non-physiological sodium concentration (25 mM) tested in the model to this point.

### Elevating Extracellular Sodium Restores Cell-Cell Electrical Connectivity Following Gap Junction Blockade

Because I_Na_, particularly in the perinexal region, is critical for ephaptic mechanisms,^21^ we hypothesized that increasing extracellular sodium might restore electrical connectivity between cardiomyocytes when GJs are functionally reduced. To test this, paired cardiomyocytes were first treated with 50-μM 2-APB for at least 30 min, and an initial recording was obtained at 25-mM extracellular sodium Na^+^. A 20-μL aliquot of concentrated Na^+^ solution (535 mM) was then added to the 1,000 μL bath (at 25 mM [Na^+^]) at a distal location in the bath, away from the recorded cell pair, raising the extracellular Na^+^ concentration to 35 mM. After a 10-s second equilibration, a second recording was acquired from the same cell pair.

We found that I_Na_ was greatly increased at 35 mM Na^+^ relative to that at 25 mM Na^+^ in the same cardiomyocyte pair (Figure 5a, top row), as reflected in the quantified I-V curves and peak I_Na_ amplitudes and densities (Figure 5a, bottom row; Figure 2Sa). More remarkably, the two-slope IDS waveform (red) reappeared at the elevated Na^+^ concentration, indicating restoration of intercellular activation. Elevating Na^+^ to 35 mM also produced a significant left-shift in steady-state activation (Figure S2b), with V_mid_ shifting from −44.31 ± 2.92 mV to −52.80 ± 4.08 mV. Because V_mid_ reflects intrinsic sodium-channel gating and is not expected to change simply with increased extracellular Na^+^,^55^ this shift is most consistent with restoration of EpC, which effectively lowers the voltage at which channels are recruited in this configuration.^20^

**Figure 5.**
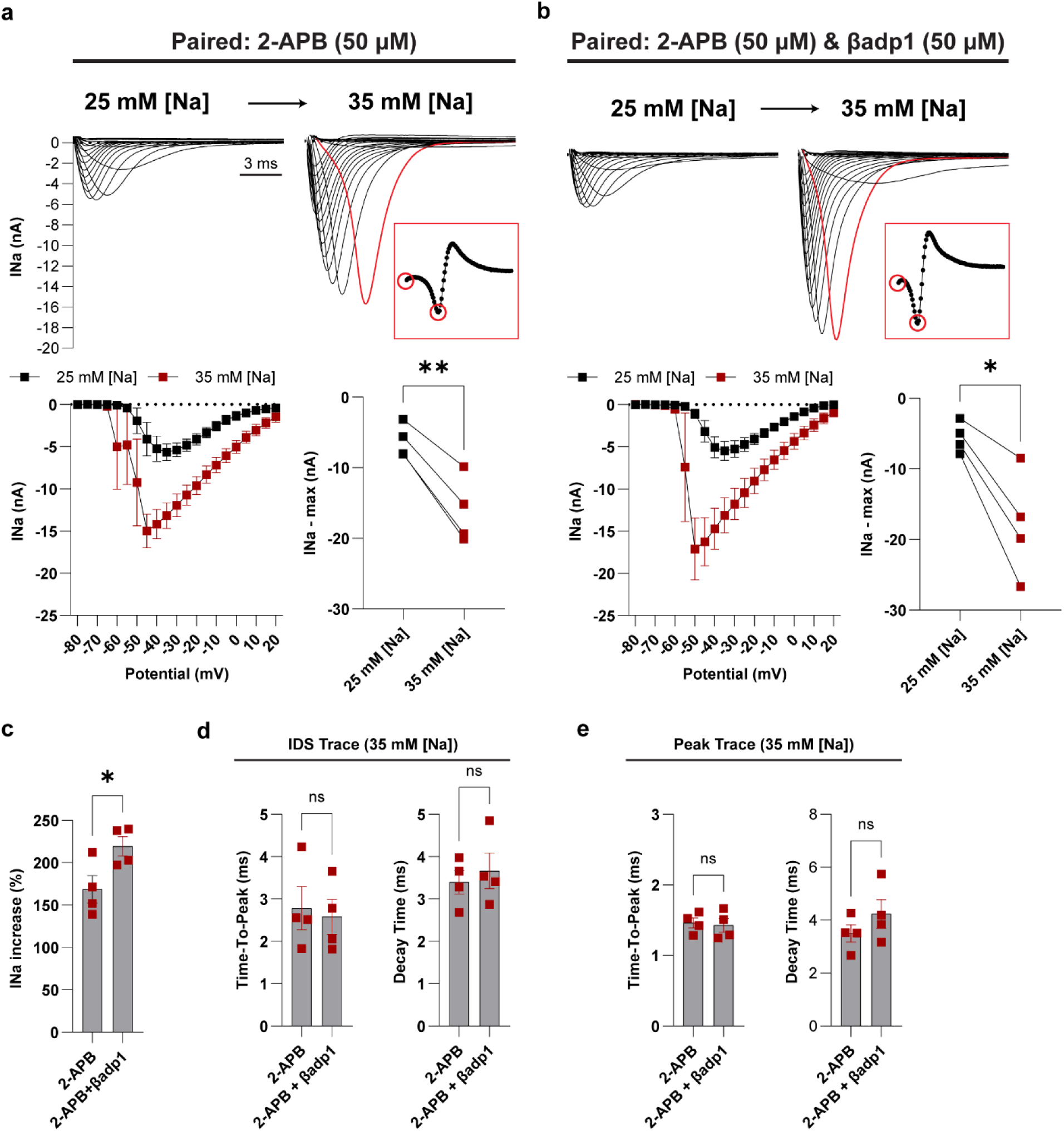
Increasing extracellular sodium restores the blocked electrical conduction between cardiomyocytes. **a**. Representative I_Na_ traces, peak I_Na_ I-V curves and max I_Na_ before and after acutely increasing extracellular sodium from 25 to 35 mM in same paired cardiomyocytes (n=4) which were treated with 50-µM 2-APB. **b**. Representative I_Na_ traces, peak I_Na_ I-V curves and max I_Na_ before and after acutely increasing extracellular sodium from 25 to 35 mM in same paired cardiomyocytes (n=4), which were treated with 50-µM 2-APB and 50-µM βadp1. **c**. The percent increase in I_Na_ from 25 to 35 mM sodium in the presence of 50-µM 2-APB, and 50-µM 2-APB & 50-µM βadp1, respectively. **d**. Time-to-peak and decay time of the IDS traces at 35-mM sodium in the presence of 50-µM 2-APB, and 50-µM 2-APB & 50-µM βadp1, respectively. **e**. Time-to-peak and decay time of the peak traces at 35-mM sodium in the presence of 50-µM 2-APB, and 50-µM 2-APB & 50-µM βadp1, respectively. Paired t-test was used for statistical analysis for panels **a** and **b**; unpaired t-test was used for panels **c**, **d**, and **e**. \**p*<0.05;\*\**p*<0.001;ns, not significant.

Using the same protocol, we next tested the effect of adding 50-μM βadp1 - inducing perinexal expansion - on I_Na_ compared with 2-APB alone. In the presence of βadp1, elevating extracellular Na^+^ again produced a robust increase in I_Na_, reappearance of the IDS waveform, and a significant left-shift of the steady-state activation curve (Figure 5b; Figure S2c and d).

To determine whether βadp1 further augments I_Na_ or facilitates trans-junctional activation beyond the effect of 2-APB alone when Na^+^ was raised from 25 to 35 mM, we quantified both the percent increase in I_Na_ and the rise/decay times of IDS and peak traces. In the presence of 2-APB, adding βadp1 significantly enhanced the I_Na_ gain (219.6 ± 11.3%) compared with 2-APB alone (168.7 ± 16%; Figure 5c). By contrast, the time-to-peak and decay times of both IDS and peak traces were not significantly altered by elevating Na^+^, regardless of βadp1 treatment (Figure 5d and 5e). These findings underscore the critical role of extracellular Na^+^ in sustaining intercellular activation when GJC is compromised and further support a perinexus-centric ephaptic mechanism, in which increased Na^+^ concentration enhances ephaptic support even when GJ conductance is strongly impaired.

### Computational Simulation of the Electrical Dynamics in Paired Cardiomyocytes

To investigate the mechanisms underlying complex whole-cell I_Na_, particularly the characteristic two-slope IDS waveform observed in the SoP model, we developed a two-cell mathematical model of electrical activity in paired cardiomyocytes based on a previously implemented model of ephaptic-mediated automaticity in cell pairs (Figure 6).^24,47^ Sodium channels from four regions were incorporated: lateral and junctional (ID) channels in both Cell 1 and Cell 2 (Figure 6, top), with ionic currents described by the Luo–Rudy 1991 formulation.^48^ The voltage-clamp protocol was applied to the intracellular node in Cell 1 (node V_1_; Figure 6, bottom). The primary goal of these simulations was to reproduce the experimental IDS and thereby examine how lateral and junctional sodium channel activation depends on extracellular sodium concentration, perinexal cleft width (W_p_) and gap junctional conductance (G_gap_). This framework allowed us to determine how different combinations of sodium concentration, GJC, and EpC shape activation of intercalated disc-adjoined cell pairs.

We found that voltage-clamping Cell 1 near the sodium channel activation threshold at −50 mV differentially affected I_Na_ in Cell 1 (solid lines) and Cell 2 (dashed lines), ultimately reproducing the IDS. Notably, this voltage step alone would recruit only minimal I_Na_ in an isolated cell (data not shown). As shown in Figure 7a, under low extracellular sodium (25 mM), nominal EpC (Wp = 10 nm) and strong GJC (G_gap_ = 700 nS), clamping Cell 1 to −50 mV recruits minimal lateral current 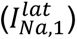, but a large junctional current 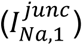 in Cell 1, and a slightly delayed but modest lateral current 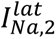 relative to 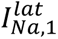) - relative to Cell 1 - and comparable junctional current (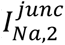 relative to 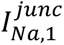) in Cell 2 (Figure 7a). Junctional sodium currents, particularly in Cell 1, clearly exhibit two slopes in the rising phase, which primarily contribute to the two-slope summed I_Na_ in the inset (pink). The junctional currents, particularly in Cell 1, clearly exhibited two slopes in the rising phase, which primarily contributed to the two-slope summed I_Na_ in the inset (pink).

**Figure 6.**
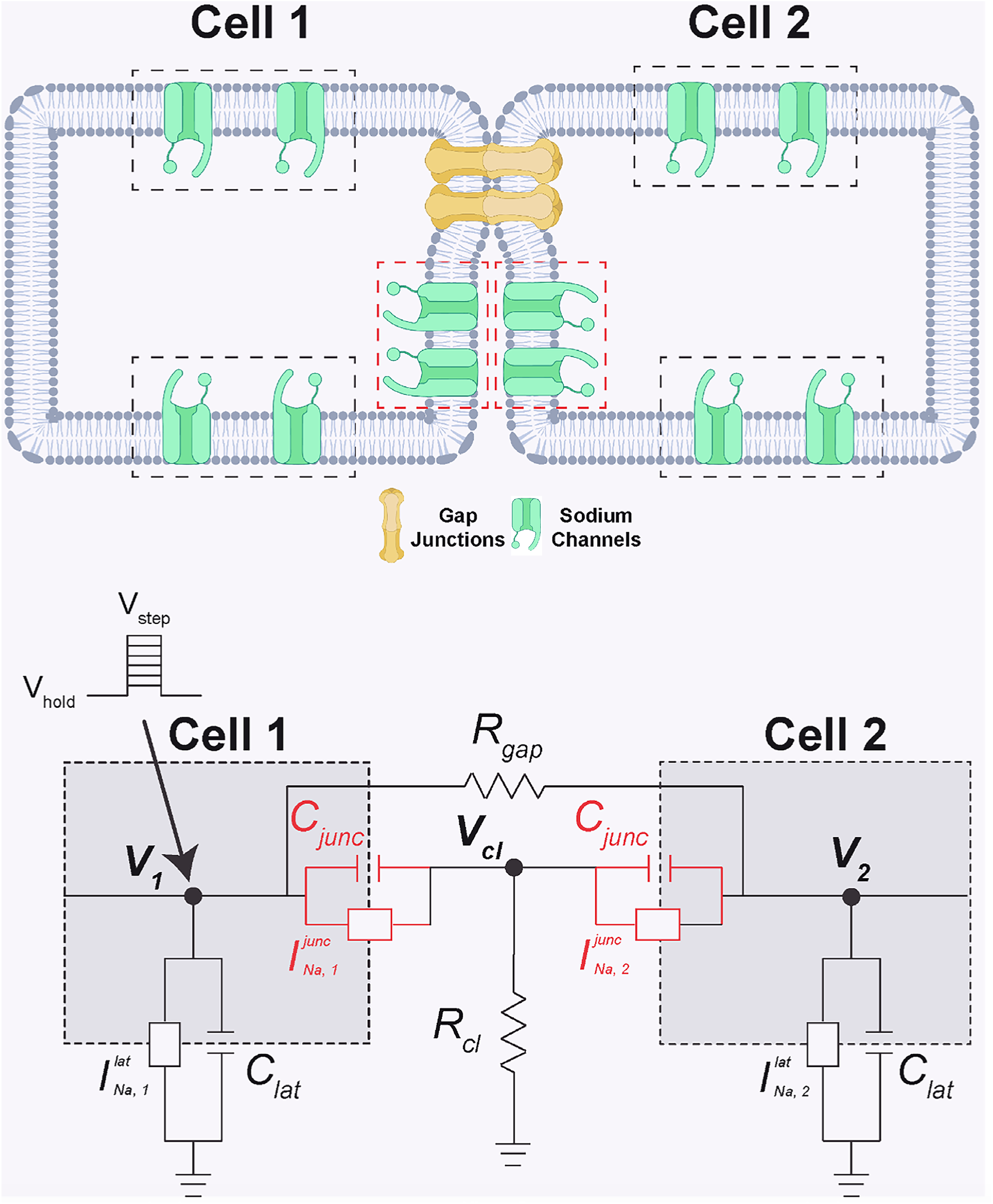
Illustration and electrical circuit for a two-cell computational model. **Top**: The illustrated two-cell model incorporates two sodium channel compartments in each cell, which were located on lateral membrane (lateral sodium channels) and junctional membrane. (junctional sodium channels-boxed region). **Bottom**: The electrical circuit of the two-cell model shows how the voltage-clamp protocol is delivered and how electrical coupling incorporating GJC and EpC is simulated.

**Figure 7.**
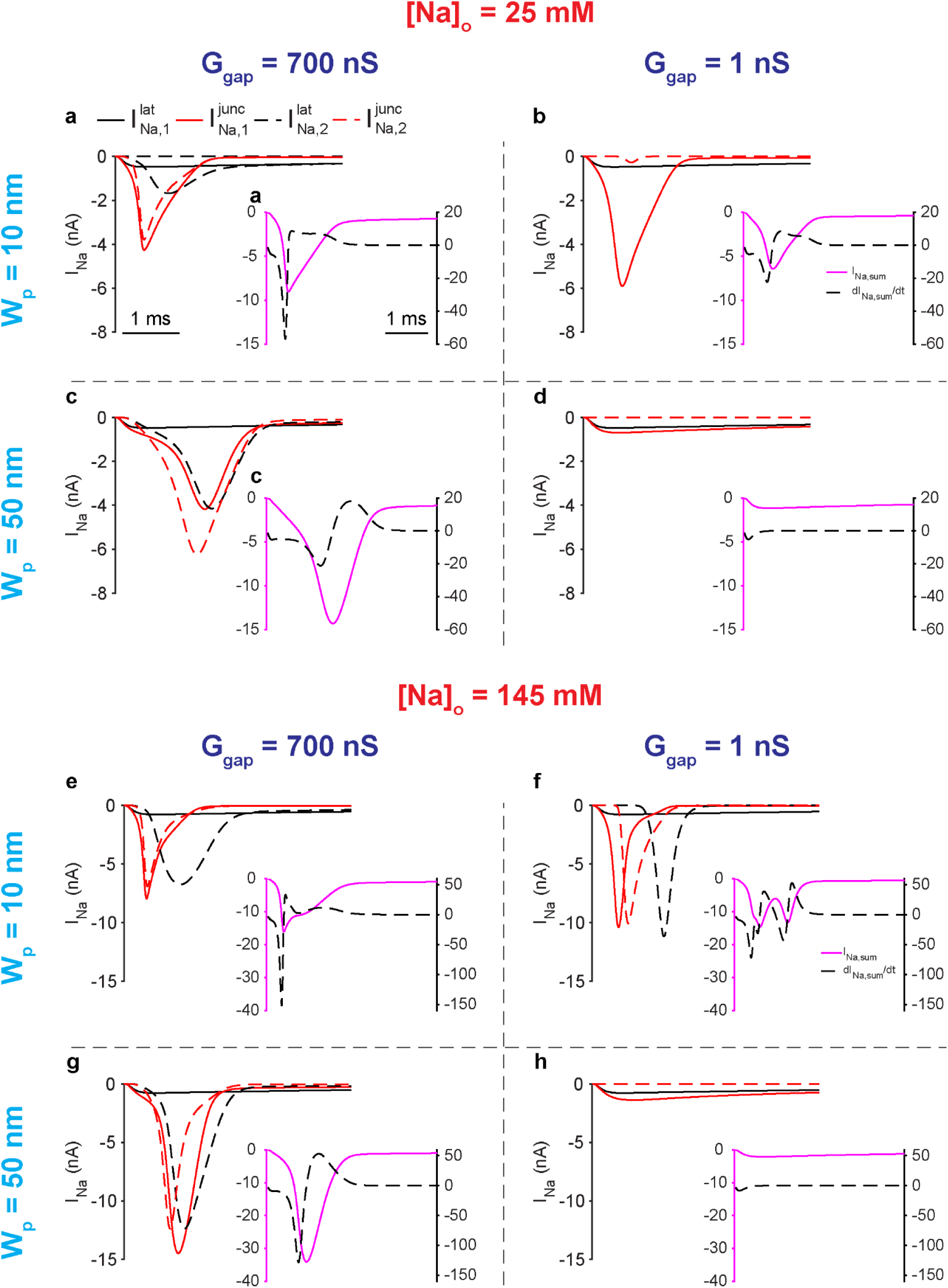
Modeling how extracellular sodium concentration shapes the interplay between gap-junctional and ephaptic coupling. **a**, **b**, **c** and **d**: the modeled responses of four compartmental sodium currents under different strengths of GJC and EpC at a voltage step of -50 mV for an extracellular sodium of 25 mM. **e**, **f**, **g** and **h**: the responses of four compartmental sodium currents under different strengths of GJC and EpC at a voltage step of -50 mV for an extracellular sodium of 145 mM.

The two-slope behavior in 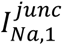 reflects an early phase dominated by the imposed voltage command at the junctional membrane and a subsequent phase in which evolving cleft potential and ephaptic interactions increasingly determine the junctional transmembrane potential, giving rise to the second slope. Clamping Cell 1 at −50 mV triggers a sufficient junctional sodium influx, driving cleft hyperpolarization (Figure S3a), which in turn increases the local transmembrane potential at the junctional membrane and thereby further increasing the junctional current, thus collectively engaging a positive feedback mechanism. Notably, this induction modifies the junctional membrane potential, relative to the imposed command voltage, such that local ephaptic interactions, rather than the clamp, dominate voltage control at the junctional membrane. The resulting more negative cleft potential, in coordination with GJC, facilitates activation of junctional sodium channels in Cell 2, further amplifying cleft hyperpolarization, followed by activation of its lateral sodium channels. Thus, the distinct timing of sodium channel activation in Cells 1 and 2 produces a summed I_Na_ with a two-slope rising phase, as evidenced by its first derivative (black dashed trace in the inset), resembling the biphasic whole-cell current recorded in the SoP model.

However, when EpC remains nominal, but GJC is low (W_p_ = 10 nm, G_gap_ = 1 nS, Na = 25 mM), only junctional sodium channels in Cell 1 are strongly activated. The cleft potential is hyperpolarized by the Cell 1 junctional sodium current (Figure S3b), however in a manner that is insufficient to activate junctional sodium current in Cell 2. Collectively, this produces a rapid, large junctional current (Figure 7b), which generates a two-slope profile in the summed whole-cell current that is now dominated by recruitment of junctional currents in Cell 1. Compared with the condition of G_gap_ = 700 nS, the lack of lateral sodium channel activation and the minimal activation of junctional channels in Cell 2 underscore the necessity of GJC for two-cell activation – at least at [25 mM Na.^+^]. These results suggest that ionic flux through GJs from Cell 1 to Cell 2 is necessary to recruit both lateral and junctional sodium channels in Cell 2 under these conditions, even in the presence of EpC.

We found that in the conditions of reduced EpC (W_p_ = 50 nm) and strong GJC (G_gap_ = 700 nS), junctional sodium channels in Cell 2 fully activate before those in Cell 1, as well as before its own lateral channels (Figure 7c). The large currents from both lateral and junctional channels in Cell 2 reflect strong GJC in the setting of a small cleft potential (Figure S3c), consistent with reduced EpC. The importance of GJC for intercellular activation at 25 mM sodium is further supported by the observation that when G_gap_ is reduced to 1 nS, currents from all regions are minimal (Figures 7d and S3d) – analogous to our observation of loss of SoP IDS with GJ inhibition at 25 mM Na^+^(Figure 5).

The simulations also showed that widening the cleft (W_p_ = 50 nm) markedly prolongs the rise time of three regional sodium currents, as reflected in the summed I_Na_ compared with a narrow cleft (W_p_ = 10 nm) when G_gap_ = 700 nS consistent with weakened EpC. In contrast, no change in the rise time of the two-slope IDS was observed following βadp1 treatment in the SoP experimental model (Figure 3). Nevertheless, the simulation reproduced the βadp1-induced increase in I_Na_ (pink traces) observed in the SoP model. Further, in the condition of G_gap_ = 1 nS, cleft widening completely abolished trans-junctional activation, again consistent with the conduction block produced by GJ inhibition in the SoP model.

Finally, we simulated activation of lateral and junctional sodium channels in both cells at physiological extracellular sodium (145 mM) while varying EpC and GJC. Under nominal EpC (W_p_ = 10 nm) and strong GJC (G_gap_ = 700 nS), activation of junctional sodium channels in Cell 1, driven by a markedly hyperpolarized cleft potential (Figure S3e), produces large currents from these channels and from all channels in Cell 2 (Figure 7e), compared with the 25 mM sodium condition (Figure 7a). When G_gap_ is reduced to 1 nS, currents from the same regions become even larger owing to the greatly decreased cleft potential (Figure S3f), reflecting dominance of EpC when GJ communication is strongly reduced (Figure 7f). Minimal GJC, however, alters the morphology of the summed I_Na_, as activation of lateral channels in Cell 2 becomes delayed relative to its junctional sodium channels.

When EpC is weakened and GJC remains strong (W_p_ = 50 nm and G_gap_ = 700 nS) at 145 mM sodium, currents from junctional channels in Cell 1 and from all channels in Cell 2 are greatly increased (Figure 7g) compared with W_p_ = 10 nm (Figure 7e), due to enhanced sodium driving force, via the smaller cleft potential (Figure S3g), a loss of so-called ‘self-attenuation,^20,21^ thus increasing sodium current in both cells under strong GJC. Strong GJC also causes junctional channels in Cell 2 to fully activate before those in Cell 1, similar to the 25-mM sodium condition with weakened EpC (Figure 7c). When both EpC and GJC are weakened (W_p_ = 50 nm and G_gap_ = 1 ns), trans-junctional activation is minimal, as shown by the small I_Na_ in both cells (Figure 7h) and the minimal cleft potential (Figure S3h). Thus, under physiological sodium levels, EpC plays a critical role in sustaining intercellular activation, especially when GJC is diminished.

Together, these simulations demonstrate that EpC and GJC mechanisms jointly and interdependently govern intercellular activation of sodium channels, with their relative contributions strongly dependent on extracellular sodium concentration. Under the non-physiologic 25 mM extracellular sodium concentration intercellular activation relies primarily on GJC, whereas from 35 mM to physiological concentrations of extracellular sodium, EpC becomes critical for maintaining electrical conduction, particularly when GJC is reduced. These results therefore indicate that extracellular sodium concentration is a key determinant of how GJC and EpC interact to mediate electrical conduction. A summary of the results and these conclusions is provided in Figure 8.

**Figure 8.**
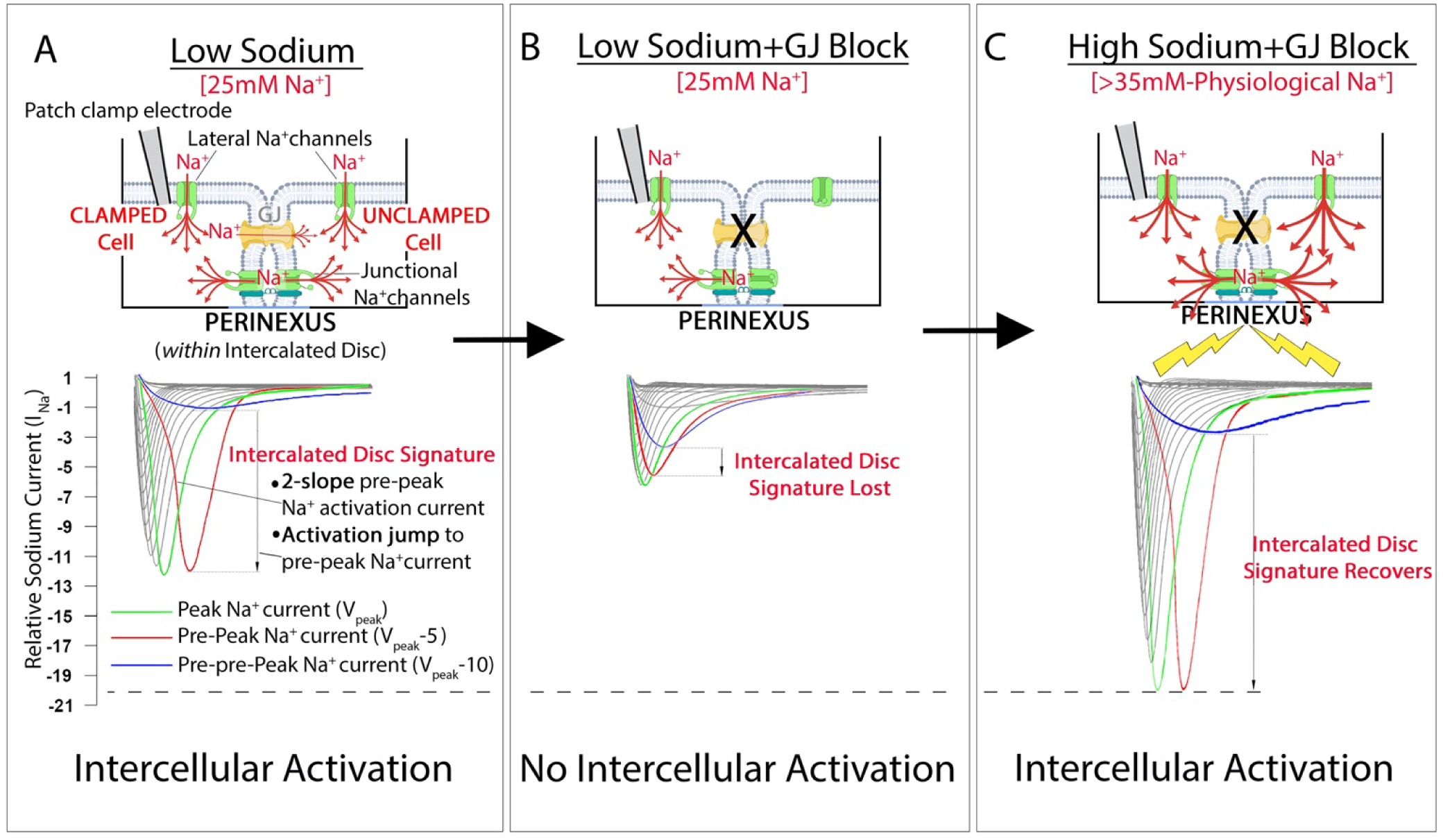
Sodium dependence of gap-junctional and ephaptic contributions to intercellular activation in the Single-on-Paired preparation. **A)** At 25 mM extracellular Na^+^ a single patch electrode clamps one myocyte that remains end-to-end coupled to an unclamped partner through an intercalated disc containing a gap junction (GJ) channel and a perinexus. Under these conditions, the whole-cell sodium current shows the Intercalated Disc Signature (IDS): a steep activation jump between closely spaced voltage steps and a two-slope rising phase in the pre-peak trace (V_peak_-5), reflecting delayed recruitment of sodium channels in the downstream cell when local cleft and downstream-cell voltages deviate from the command voltage, with simulations indicating that junctional (perinexal) channels are the primary contributors to this delayed component relative to lateral channels. **C)** When GJ coupling is strongly inhibited at the same low Na^+^, the IDS disappears and peak I_Na_ falls to approximately single-cell levels, indicating loss of detectable intercellular activation; in this low-sodium regime, ephaptic interactions alone are insufficient to sustain trans-junctional activation. **C)** Elevating extracellular [Na^+^] to 35 mM in the continued presence of gap-junction block restores the IDS and large whole-cell currents. Together with the two-cell simulations at physiological Na^+^, this indicates that intercellular activation is then supported predominantly by perinexus-centered ephaptic mechanisms acting through the same Intercalated Disc nanodomain. Overall, the data show that the balance between GJC and EpC is strongly sodium-dependent. GJC dominates at non-physiologic low 25 mM Na^+^, whereas as Na^+^ approach more physiologic levels, ephaptic coupling can provide robust support for activation, even when GJC is compromised. Moreover, the IDS provides an operational readout of how the *gap junction–perinexus nanodomain* integrates junctional conductance, cleft geometry, and sodium concentration to govern cardiomyocyte-to-cardiomyocyte electrical communication.

## DISCUSSION

In this study, we introduce a ‘Single-on-Paired’ (SoP) cardiomyocyte preparation that isolates gap junctional coupling (GJC) and ephaptic coupling (EpC) by resolving a distinct ‘Intercalated Disc Signature’ (IDS) within the whole-cell sodium current (I_Na_) of myocyte pairs. The IDS comprises two linked features: a steep activation “jump” in I_Na_ between closely spaced voltage steps and a characteristic two-slope rising phase that appears selectively at the pre-peak trace in paired myocytes. Together with a complementary two-cell computational model, these data show that GJC alone can sustain intercellular activation at 25 mM Na^+^, whereas EpC becomes a critical contributor as extracellular Na^+^ approaches physiological level. We therefore formalize a previously implicit concept: the *gap junction–perinexus nanodomain* as a governing determinant of cardiac electrical conduction.

The SoP model preserves a native intercalated disc (ID) between two myocytes while retaining the technical simplicity of a single whole-cell patch-clamp configuration, allowing I_Na_ to report intercellular activation in real time. Our data show that paired cells maintain native arrangements of ion channels at key domains, including gap junctions (GJs) and the perinexus,^31,56^ structures that herein we, and others previously,^53^ have shown are internalized and largely lost in isolated single myocytes. In this context, paired myocytes exhibit a distinctive IDS that appears only when an intact ID connects the clamped and downstream cell and is selectively modulated by GJ inhibition and perinexal remodeling. Unlike conventional single-cell recordings, which are restricted to local ionic currents, or dual patch-clamp of cell pairs, which is technically demanding and primarily used to quantify GJ conductance,^57^ the SoP configuration combines single-patch practicality with the ability to probe the interplay between GJC and EpC, providing a platform for future studies of intercellular action potential propagation and activation of additional channel populations regardless of cell type.

The main innovation in this study is the discovery of the IDS, which raises a reasonable question: Is the IDS a technical artifact without biological relevance or a robust reporter of GJC and EpC? While we never observed a two-slope I_Na_ trace in isolated single myocytes, others have reported such behavior in overexpressing systems under conditions of clear loss of voltage control due to insufficient series resistance compensation or ‘space clamp’.^58–65^ In contrast, in our SoP preparation the IDS never appears in isolated cells, is abolished by gap junction uncoupling at 25 mM extracellular Na^+^ and reappears in the same GJ-inhibited cell pair when Na^+^ is raised to 35 mM. These observations argue that imperfect series resistance compensation alone is unlikely to account for these manifestations of the IDS. We recognize that any whole-cell experiment is subject to space-clamp limitations; accordingly, we also examined large single cardiomyocytes, in which such issues are most likely to be evident. Again, we found no IDS like waveforms in single cells, whereas bona fide cardiomyocyte pairs consistently exhibited this signature.

Voltage clamping also assumes that the transmembrane potential at the pipette tip is uniform throughout the clamped cell (i.e., the cell is isopotential). This approximation holds when intracellular and axial resistances are low, but GJs introduce much higher resistance than the myocyte interior. In the SoP configuration, the patched cell (Cell 1) is therefore reasonably space clamped, whereas the downstream partner (Cell 2) is not. Our SoP methodology deliberately exploits this controlled loss of voltage control in Cell 2: instead of using voltage clamp solely to measure peak sodium current in a single cell, we use it to distinguish GJC from EpC in a two-cell system by analyzing timing. Under conditions of very low extracellular sodium, and at test potentials that already produce somewhat dyssynchronous sodium channel recruitment in isolated cells, the same command step applied to Cell 1 in the SoP preparation first elicits a small, desynchronized I_Na_ in the clamped cell, and then, after a delay, an unclamped I_Na_ transient arising from Cell 2, with quantifiably distinctive biophysical properties.

A related HEK–obstacle preparation developed by Hichri and colleagues provided independent support for our findings. By moving a sodium channel transfected HEK cell toward an inert barrier to create an artificial perinexal-like cleft, they showed that narrowing cleft volume alone can markedly reshape I_Na_, consistent with strong ephaptic modulation of I_Na_ in the absence of myocyte-specific architecture. In this non-myocyte system, the loss of voltage control occurs extracellularly at the interface of the myocyte and inert barrier. Their non-myocyte system therefore converges with our SoP data and simulations in indicating that local cleft geometry and resistive coupling with a second cell are sufficient to generate two-slope, field-sensitive I_Na_ waveforms that report trans-junctional activation.

Taken together, the kinetics associated with the controlled loss of voltage control in the SoP configuration enable us to distinguish GJC- and EpC- mediated contributions to intercellular activation. Specifically, experimental and computational data show that in this model: (i) conduction can be maintained by GJC alone at low extracellular sodium [25 mM] ; (ii) widening the perinexus modulates ephaptic strength and can augment the summed sodium current in paired cells; (iii) strong GJ inhibition abolishes intercellular activation at low sodium; and (iv) a modest 10 mM increase in extracellular sodium [from Na^+^ 25 mM] during GJ inhibition is sufficient to restore the IDS via perinexus-centered EpC. Together, these findings support the view that ephaptic communication is a major determinant of electrical communication between cardiomyocytes at physiologic sodium concentrations, and that the IDS provides a robust operational readout of whether intercellular activation in the SoP preparation is dominated by GJC, EpC, or their interaction.

The SoP protocol also clarifies whether GJC and EpC act as independent or cooperative mechanisms. Classic Cx43 loss-of-function studies show that ventricular conduction can persist despite substantial reductions in GJC, leading to the idea that ephaptic mechanisms maintain propagation when GJ channels are compromised.^42,66,67^ In the SoP model, nominal GJ inhibition abolishes detectable intercellular activation in the presence of an intact perinexus, implying that ephaptic support of conduction requires a baseline level of GJC, at least under the non-physiologic [25 mM] sodium condition. When extracellular sodium is increased, however, the IDS reappears and perinexal widening enhances conduction despite persistent GJ inhibition, indicating that with more physiological sodium concentrations, EpC can maintain intercellular activation. Thus, rather than acting as mutually exclusive pathways, GJC and EpC can behave as dynamically reconfigurable components of a shared conduction apparatus whose relative contributions are tuned by sodium gradients and nanoscale cleft geometry.

Our simulations help generalize these ideas and place them in the context of prior modeling work.^20,22,37^ By explicitly resolving lateral versus junctional sodium-channel populations and including both GJC and a resistive intercellular cleft, the two-cell model reproduces the IDS and recapitulates transitions in the relative contributions of GJC and EpC as extracellular sodium and cleft geometry are varied. At physiologic sodium concentrations, the model predicts that reduced GJC can be offset by strong perinexal ephaptic interactions, allowing opposing junctional sodium-channel clusters across a narrowed cleft to sustain intercellular activation, even when direct coupling is limited. In contrast, under non-physiologic, markedly lowered sodium, conduction in the model becomes increasingly dependent on GJC and ultimately fails when GJC is strongly reduced, consistent with the SoP experiments at 25 mM Na⁺. Notably, the timing effects of perinexal widening on the IDS still differ in detail between experiment and our current model, underscoring that the precise temporal interplay of GJC and EpC in IDS generation remains incompletely defined and will require further refinement and validation in two-cell models.

The SoP experiments were necessarily performed at reduced extracellular sodium (25–35 mM), because rapid sodium channel activation in isolated myocytes cannot be reliably resolved under voltage clamp at fully physiological sodium levels. Even so, combining SoP data with the two-cell computational model allowed us to interrogate the GJC–EpC balance across a wider sodium range and perinexal widths, including physiological conditions. Both experimental and simulation results indicate that extracellular sodium is a key determinant of whether GJC alone can maintain conduction or whether EpC must be recruited, reinforcing our previous findings that sodium concentrations can in part explain why whole-heart Cx43 knockouts often show preserved or only modestly reduced conduction at baseline.^68^

Although the perinexus occupies only a 10–30 nm wide intercellular cleft, dense Nav1.5 clustering means that a non-trivial fraction of whole-cell I_Na_ is injected into an extracellular annulus on the order of tens of nanometers surrounding each Cx43 plaque,^32,56,69,70^ particularly when sodium channel clusters face each other across the cleft, as we and other have proposed.^18,71^ Modeling and ultrastructural studies suggest that such arrangements yield effective current densities on the order of tens of mA/cm² and local electric fields approaching hundreds to thousands of V/m during the IDS upstroke^22,37,72^ - hence the lightning bolts on panel C of the summary model (Figure 8). In this respect, the perinexal cleft behaves less like a diffuse extracellular space and more like the nanometer-scale gap at a transistor gate, where small, highly localized changes in charge and voltage are sufficient to switch an entire circuit between off and on states. Viewed this way, the IDS reflects a surprisingly strong, tightly focused, sodium-driven field effect, reinforcing the idea that at physiological extracellular Na⁺, the perinexus-centered ephaptic mechanism is a potent driver of intercellular action potential transfer rather than a minor adjunct to GJC, as we have posited in an earlier study. ^19^

Several limitations temper extrapolation of these findings to intact organs and other tissues. The SoP model uses isolated ventricular myocyte pairs that inevitably undergo some ID remodeling, and recordings were restricted to sub-physiological sodium levels. Electrical block was induced pharmacologically rather than by genetic ablation of Cx43, and 2-APB is an incompletely selective and incompletely blocking inhibitor whose effects on GJ conductance and intracellular signaling may be complex.^50^ This being said, studies of the ephaptic effects of single Purkinje neurons also reported the use of 2-APB as more favorable than other pharmacologic GJ inhibitors such as carbenoxolone and 18β-glycyrrhetinic acid,^12^ both of which we found toxic on isolated cardiomyocytes. Our SoP configuration also cannot directly record downstream I_Na_ when junctional resistance is very high, so residual ephaptic currents below detection cannot be excluded under strong GJ inhibition. Future use of cardiomyocyte-specific Cx43 knockout or more selective GJ modulators, together with refined modeling of IDS timing, will help define the precise GJC dependence of EpC in this preparation.

Despite these caveats, the present work positions the SoP model as a versatile platform for dissecting how Cx43-anchored nanodomains integrate GJC and EpC to regulate activation. By identifying extracellular sodium and cleft nanostructure as key determinants of the balance between direct and field-mediated coupling, and by demonstrating that a perinexus-centered ephaptic mechanism can provide conduction reserve when GJC is compromised, this study refines our understanding of ventricular conduction and suggests the potential for mechanistically informed strategies for targeting sodium channels, β-subunits, or perinexal structure in arrhythmia therapy. Similar Cx43-anchored nanodomains in astrocyte syncytia^73^ and in the myocardium of hibernating mammals,^74^ where electrical activity is preserved despite profound ionic and metabolic remodeling, may reflect parallel strategies for dynamically rebalancing GJC and EpC to maintain circuit function under stress. More broadly, these findings raise testable hypotheses for these and other Cx43-rich tissues - including His-Purkinje fibers,^75^ and myometrium at term^76^ - in which coordinated variation of GJC and EpC may shape the efficacy and failure of anti-arrhythmic and anti-epileptic interventions in normal and diseased states.

## ACKNOWLEDGMENTS

The authors thank Dr. Matthew Weston from Fralin Biomedical Research Institute at Virginia Tech and Dr. Xianming Lin from New York University Langone Health for their valuable technical advice during this study.

