## Supplemental figures for "Dissecting Gap Junctional and Ephaptic Contributions to Electrical Conduction in a Novel Cardiomyocyte Pair Model"

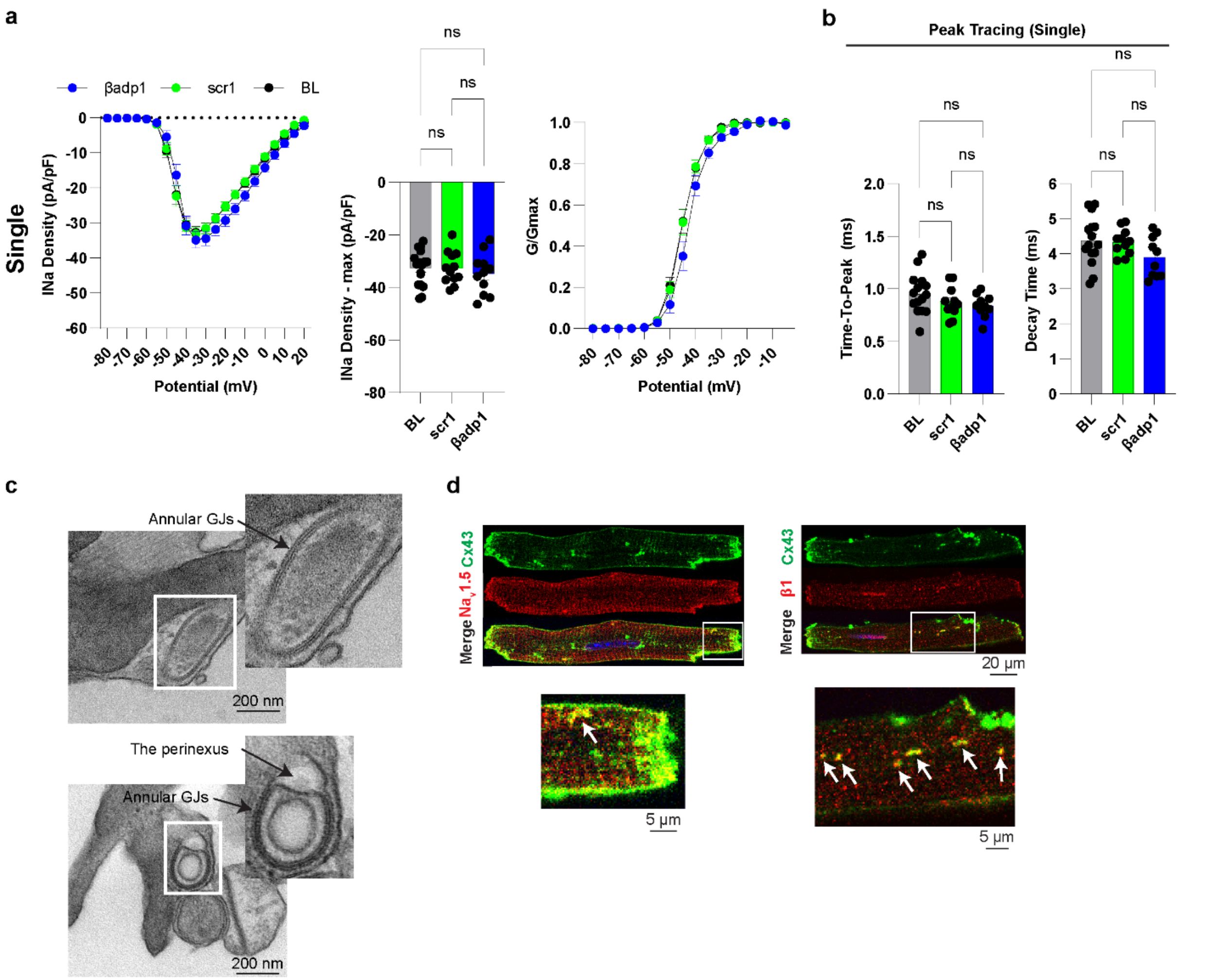


**Figure S1. Perinexal remodeling increases sodium current in paired cardiomyocytes (continued). a**. INa density I-V curves, max INa densities and steady-state activation curves in single cardiomyocytes at BL (n=17), scr1 (n=13) and βadp1 (n=11). **b**. Time-to-peak and decay time of the peak trace in single cardiomyocytes at BL, scr1 and βadp1. **c**. The internalization of gap junctions and perinexus near IDs in single cardiomyocytes. **d**. Immunofluorescent signals of Cx43&Nav1.5, Cx43&β1/β1b inside single cardiomyocytes. One-way ANOVA followed by Tukey correction for multiple comparison for **a** and **b**. *$p<0.05$; ns, not significant.

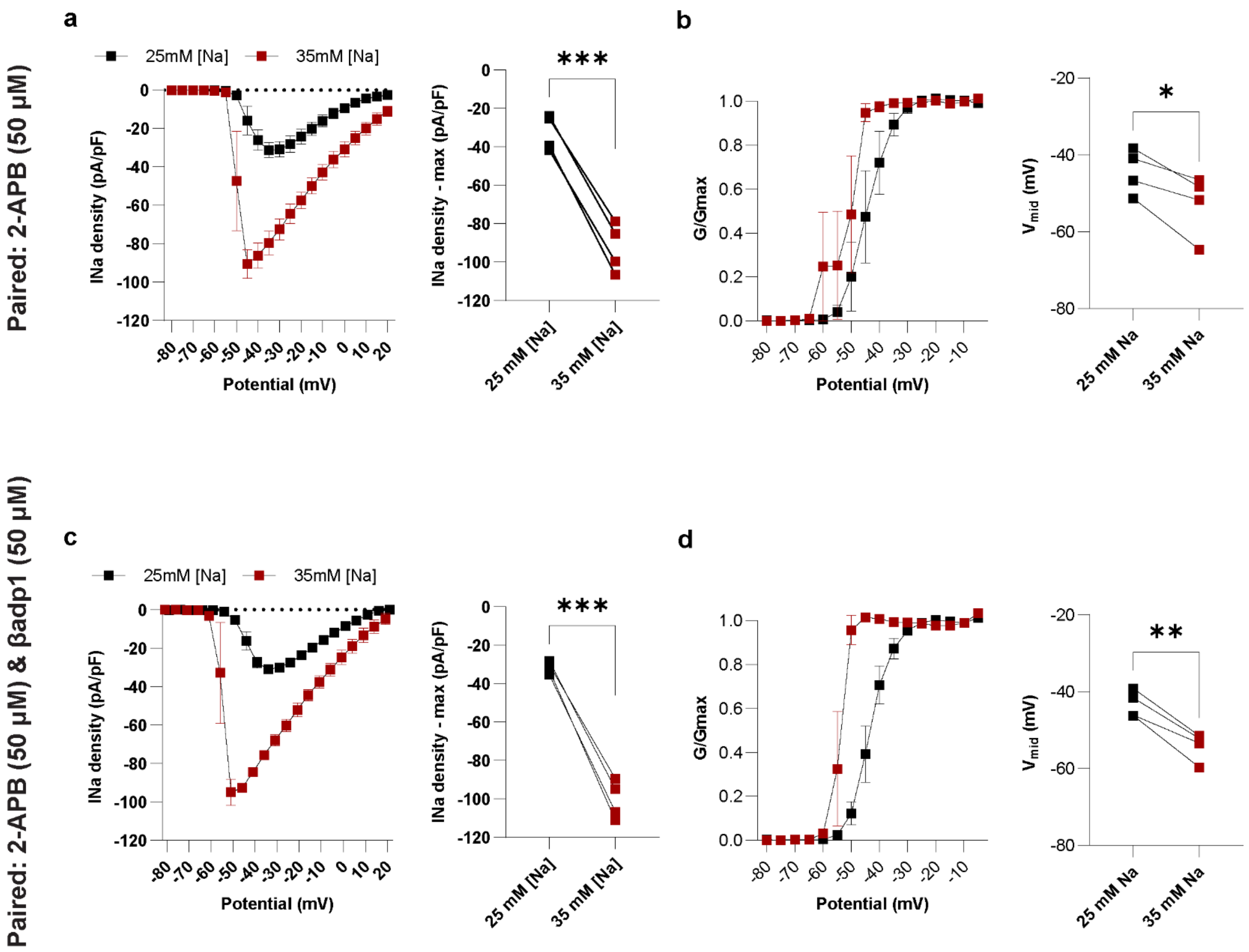
Paired: 2-APB (50 µM)
a
25mM [Na]
35mM [Na]
INa density (pA/pF)
Potential (mV)
***
INa density - max (pA/pF)
25 mM [Na]
35 mM [Na]
b
G/Gmax
Potential (mV)
*
$V_{mid}$ (mV)
25 mM Na
35 mM Na
Paired: 2-APB (50 µM) & βadp1 (50 µM)
c
25mM [Na]
35mM [Na]
INa density (pA/pF)
Potential (mV)
***
INa density - max (pA/pF)
25 mM [Na]
35 mM [Na]
d
G/Gmax
Potential (mV)
**
$V_{mid}$ (mV)
25 mM Na
35 mM Na

**Figure S2. Increasing extracellular sodium restores the blocked electrical conduction between cardiomyocytes (*continued*). a**. $I_{Na}$ density I-V curves and max $I_{Na}$ densities before and after acutely increasing extracellular sodium from 25 to 35 mM in same paired cardiomyocytes (n=4) which were treated with 50-μM 2-APB. **b**. Steady-state activation curves and $V_{mid}$ before and after acutely increasing extracellular sodium from 25 to 35 mM in same paired cardiomyocytes (n=4) which were treated with 50-μM 2-APB. **c**. $I_{Na}$ density I-V curves and max $I_{Na}$ densities before and after acutely increasing extracellular sodium from 25 to 35 mM in same paired cardiomyocytes (n=4) which were treated with 50-μM 2-APB and 50-μM βadp1. **d**. Steady-state activation curves and $V_{mid}$ before and after acutely increasing extracellular sodium from 25 to 35 mM in same paired cardiomyocytes (n=4) which were treated with 50-μM 2-APB and 50-μM βadp1.

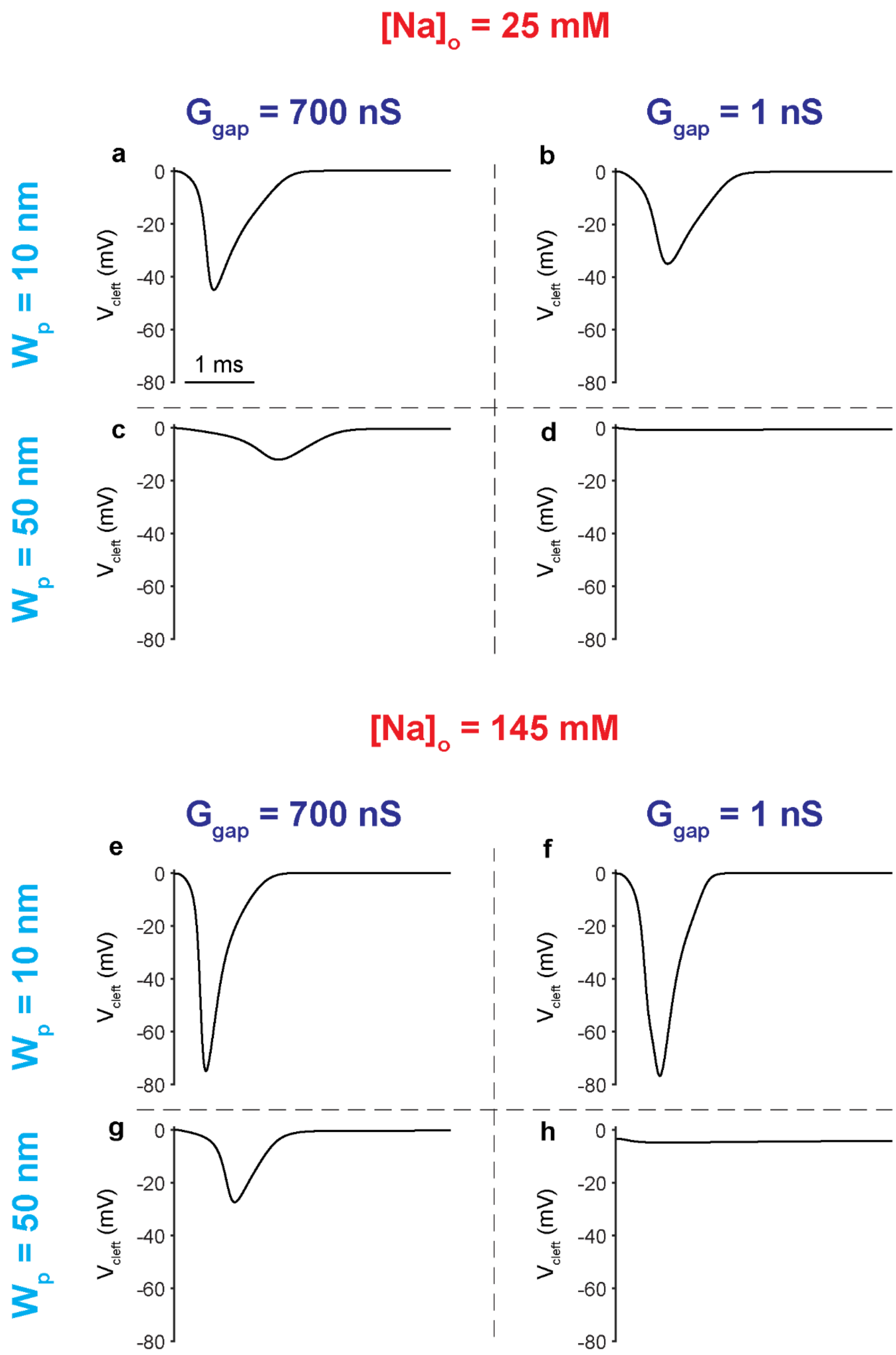

[Na]$_o$ = 25 mM
G$_{gap}$ = 700 nS
G$_{gap}$ = 1 nS
W$_p$ = 10 nm
W$_p$ = 50 nm
a
b
c
d
V$_{cleft}$ (mV)
0
-20
-40
-60
-80
1 ms
[Na]$_o$ = 145 mM
G$_{gap}$ = 700 nS
G$_{gap}$ = 1 nS
W$_p$ = 10 nm
W$_p$ = 50 nm
e
f
g
h

**Figure S3. Extracellular sodium concentrations determine the interplay between gap junctional and ephaptic coupling (*continued*).** **a**, **b**, **c** and **d**: Cleft potentials under different strengths of GJC and EpC at a voltage step of -50 mV when the extracellular sodium is 25 mM. **e**, **f**, **g** and **h**: Cleft potentials under different strengths of GJC and EpC at a voltage step of -50 mV when the extracellular sodium is 145 mM.
